# Resolving symmetry-masked allosteric cooperativity in the *M. tuberculosis* proteasome core particle

**DOI:** 10.64898/2026.07.02.736129

**Authors:** Madison Turner, Verena Wittlinger, Cristina Lento, Anisha Haris, Jakub Ujma, David Bruton, Keith Richardson, Kevin Giles, Thomas Böttcher, Derek J. Wilson, Siavash Vahidi

## Abstract

The 20S proteasome core particle (CP) is a stacked α_7_-β_7_-β_7_-α_7_ assembly in which the central β-rings host fourteen catalytic active sites responsible for regulated protein degradation. Allosteric coupling between catalytic β-subunits has been characterized in eukaryotic proteasomes, whose heteromeric β-rings permit subunit-specific perturbation. In bacterial proteasomes, however, the β-rings are homomeric, and any allosteric relationships between subunits with identical sequences have remained refractory to conventional ensemble-averaging structural methods, including to hydrogen/deuterium exchange mass spectrometry (HDX-MS). Here we show that orthosteric inhibitors paradoxically activate the *Mycobacterium tuberculosis* 20S CP at substoichiometric concentrations, revealing positive cooperativity between its β-subunits. To dissect this cooperativity within the β-ring, we co-assemble wild-type and catalytically inactive (T1A) β-subunits into hybrid 20S CPs. We develop a probabilistic model relating bulk mixing ratios to the ensemble of hybrid 20S CP stoichiometries. Differential ^15^N-labelling of the wild-type subunits then resolves WT and T1A peptide signals by mass during HDX-MS, enabling subunit-resolved measurements within a single complex. Using this approach, we demonstrate that ligand binding at one β-subunit remodels the conformational dynamics of binding-incompetent neighbours. Measuring deuterium uptake against ring composition identifies two allosteric routes: a lateral pathway from switch helix II to the active site of the adjacent intra-ring subunit, and an axial pathway connecting a loop at the β-ring interface to the S pockets of the opposing ring. More broadly, this work establishes a framework for resolving symmetry-masked allostery in multi-subunit assemblies.

**Significance Statement:** The 20S proteasome is essential for the survival and virulence of *Mycobacterium tuberculosis*, yet how its catalytic sites communicate has been difficult to study because the bacterial enzyme is built from identical subunits whose signals are indistinguishable by conventional methods. Here we find that blocking only a fraction of these sites by inhibitors paradoxically activates the enzyme, revealing positive cooperativity between neighbouring subunits. To trace the communication pathways, we developed an isotopic-coding strategy that enables hydrogen/deuterium exchange mass spectrometry report on individual subunits within a single symmetric complex. This approach maps how ligand binding at one subunit reshapes its neighbours and, more broadly, provides a general framework for dissecting allostery in homomeric molecular machines whose symmetry has long obscured it.

## Introduction

Maintaining cellular homeostasis requires the degradation of damaged or redundant proteins. This necessitates a highly targeted, specific process such that only intended protein substrates are proteolyzed under appropriate cellular conditions (1–3). Proteasome systems in both eukaryotic and prokaryotic organisms form bipartite assemblies in which regulatory particles (RPs) recognize specific substrates and deliver them to the 20S core particle (CP) for proteolysis (4–7). The 20S CP is a barrel-shaped oligomer formed by the stacking of four heptameric rings in a conserved α_7_-β_7_-β_7_-α_7_ configuration (Fig. 1A) (8–12). Catalytic sites within the β-subunits are sequestered within the lumen of the degradation chamber (Fig. 1B), while the α-rings regulate access to this chamber through conformational gating (13, 14). RP binding to the α-rings coordinates substrate engagement, gate opening, and translocation into the proteolytic chamber (15–18). These layers of regulation are further augmented by allostery embedded within the proteasome architecture (19–21).

**Figure 1.**
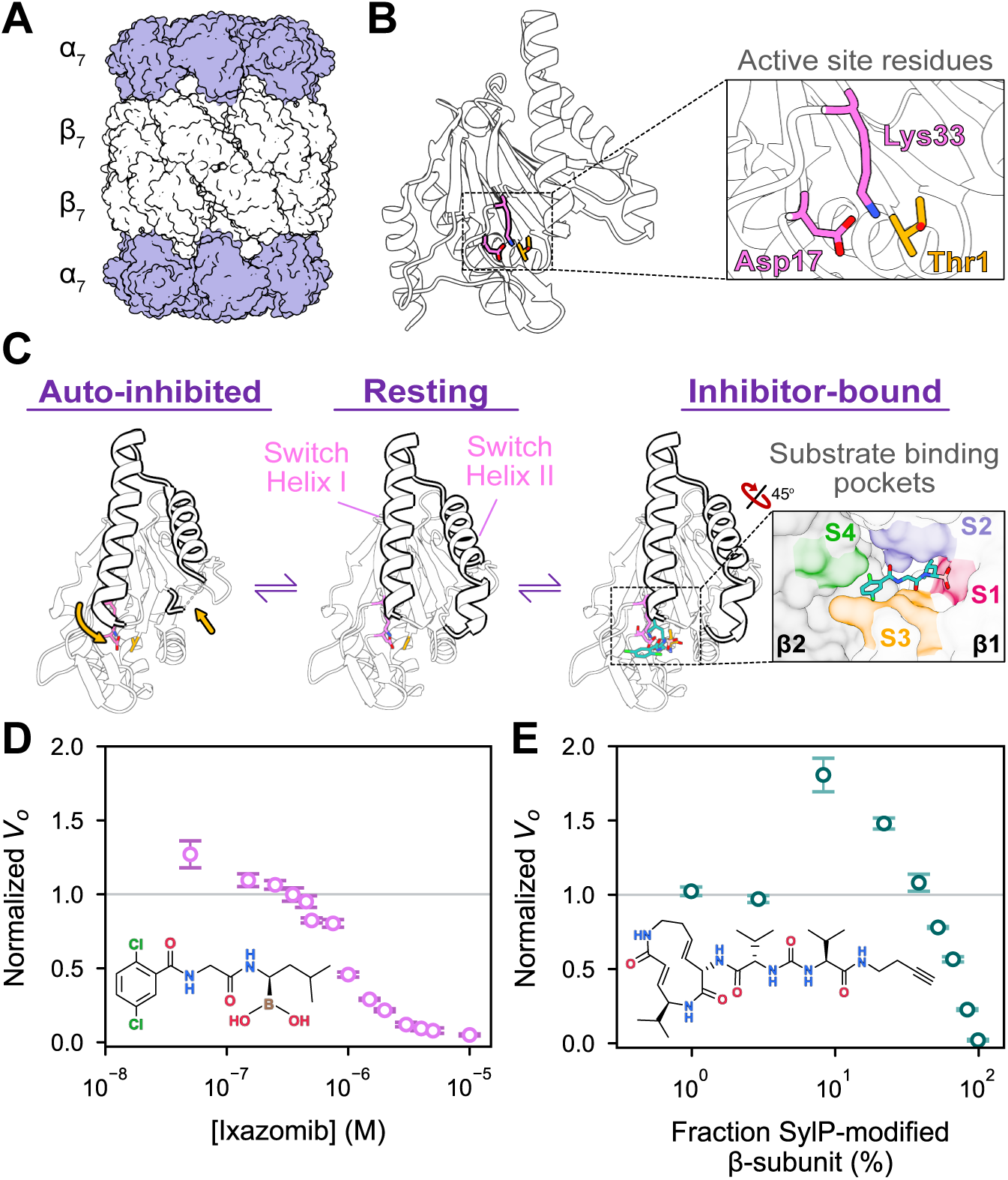
The *Mtb* 20S CP is an allosteric enzyme with varying β-subunit conformations. (A) The *Mtb* 20S CP is formed through the staking of four homo-heptameric rings in an α_7_-β_7_-β_7_-α_7_ configuration (PDB 9ce5); (B) Substrate degradation is carried out by the β-subunits via a catalytic triad formed through Thr1, Asp17 and Lys33 (PDB 9ce5); (C) The conformation of the switch helices modulates access to the β-subunit active sites. Conversion between the auto-inhibited state (PDB 9cee) and the resting state (PDB 9ce5) of the protein requires a shift in switch helix I upon restructuring of the C terminus of switch helix II (changes are denoted by yellow arrows, left). This conformational change enables the engagement of ligand within the active site, which is comprised of four distinct substrate-binding pockets (S1-4, right; PDB 9ce8); and (D, E) The binding of covalent orthosteric inhibitors at sub-stoichiometric concentrations activates the peptidase function of the *Mtb* 20S CP.

Allostery enables regulation of protein function in response to effector binding (22–24). Ligand binding redistributes conformational populations within the protein, thereby modulating interactions with ligands or binding partners and tuning activity (22–24). In multi-subunit assemblies, these conformational changes give rise to cooperativity, whereby local binding events influence distal functional sites within the same subunit or across neighbouring subunits. Extensive evidence supports cooperative allostery across the α/β-interface in proteasome systems. For example, the ligation state of β-subunits can modulate the affinity of RP engagement at the α-ring (25–28). In addition, binding of a first RP to one α-ring can alter the affinity of a second RP at the distal α-ring, demonstrating long-range cooperativity across the entire length of the 20S CP (20, 29, 30). In contrast, direct cooperativity between the two opposing β-rings appears weaker, with archaeal studies reporting minimal β:β cross-ring coupling in the absence of regulatory particles (29). Likewise, β-subunit catalytic activity responds to RP binding (31, 32) and to conformational changes within the α-subunit (33, 34). In the eukaryotic 20S CP, the β-ring is heteromeric, each of its seven β-subunits is chemically distinct, and lateral cooperativity between β-subunits has been characterized using the ligation state of one subunit to perturb its neighbours (35–37). The prokaryotic 20S CP, by contrast, contains a homomeric β-ring whose seven identical subunits render the same allostery refractory to ensemble study, because each subunit contributes an indistinguishable signal.

Although high-resolution structural studies have defined proteasome architecture in detail, comparisons of available structures often reveal only modest conformational differences, even in the presence of ligands or regulatory particles (14, 38–40). This suggests that functionally relevant allostery in proteasomes is, at least in part, encoded by shifts in conformational dynamics rather than in large-scale architectural rearrangements. Accordingly, solution-phase approaches such as NMR spectroscopy and H/D exchange mass spectrometry (HDX-MS) have provided complementary insight into these dynamic changes and the cooperative mechanisms that underlie them (25, 27, 29, 33, 41). To interrogate cooperativity in a symmetry-constrained system, we recently investigated the bacterial 20S CP from *Mycobacterium tuberculosis* (33). We demonstrated that a pair of β-subunit helices at the α/β-interface, switch helix I and II, adopted a unique conformation different from the conformation seen in the canonical structure (33). Localized unwinding of the C terminus of switch helix II initiated a positional shift in switch helix I, thereby collapsing part of the substrate binding pocket (S1 pocket) (Fig. 1C). We showed that this auto-inhibited state of the 20S CP interconverts with the canonical resting state capable of engaging substrate. Moreover, targeting allosteric sites up to 48 Å away on the α-subunit modulates this equilibrium, demonstrating long-range cooperativity across the α/β-interface (33). Given that the switch helices span both the α-β interface and the β-β interface while contributing directly to the substrate-binding pocket, conformational changes at these elements are ideally positioned for lateral allosteric coupling to the neighbouring β-subunits. In the symmetric *Mtb* 20S CP, however, the sequence identity of all β-subunits obscures the detection of such communication by conventional structural approaches.

Here, we uncover a previously unrecognized mode of allosteric modulation within the *Mtb* 20S CP. We discovered that orthosteric inhibitors, classically defined by their ability to suppress catalysis, paradoxically activate the peptidase function of the 20S CP at substoichiometric concentrations. This counterintuitive behaviour suggests that engagement of only a subset of β-subunits is sufficient to promote activation of the entire complex, and is consistent with inter-β ring cooperativity within the 20S CP. Resolving the structural basis of this phenomenon presents a fundamental challenge: in symmetric complexes with partial ligand occupancy, ensemble-averaging structural approaches struggle to disentangle heterogeneous ligation states or to resolve responses among identical subunits. In conventional HDX-MS, this degeneracy masks whether conformational changes arise from direct effector binding or from allosteric propagation to neighbouring subunits. By integrating stable isotopic coding with HDX-MS, we break β-subunit degeneracy and achieve subunit-resolved analysis of conformational dynamics within the 20S CP. Using this approach, we demonstrate that ligand engagement at one β-subunit active site is associated with coordinated conformational changes in neighbouring β-subunits, implicating the switch helices and surrounding regions in mediating allosteric coupling across the β-ring and defining a structural pathway for cooperativity within the symmetric protease chamber. More broadly, this work defines a general framework for resolving symmetry-masked allostery in multi-subunit complexes.

## Results

### Allosteric activation of the *Mtb* 20S CP via substrate-mimic binding

Peptidyl boronates have previously been shown to inhibit *Mtb* 20S CP peptidase activity by engaging the nucleophilic βThr1 via their boron warhead (13, 33), thereby blocking catalytic turnover (42, 43). Strikingly, however, characterization of the peptidyl boronate ixazomib revealed a paradoxical activation of enzymatic activity at substoichiometric concentrations (Fig. 1D). At [ixazomib] below 0.35 µM, peptidase activity increased up to 30% relative to the apo control, while higher concentrations displayed the expected competitive inhibition (Fig. 1D). To ascertain whether this effect was specific to the boron warhead, we next examined the syrbactin derivative, syringolin A (SylA), a mechanistically distinct inhibitor which has previously been used to target the eukaryotic 20S CP and immunoproteasome (44). Like other syrbactins, SylA targets the catalytic Thr1 residue by forming a covalent linkage via a Michael-type addition (44, 45). Unlike the covalent bond formed by peptidyl boronates, this linkage is non-hydrolyzable resulting in an irreversible inhibition. We used a chemically modified syringolin A probe (SylP) bearing a terminal alkyne handle, which promotes access to the proteolytic core chamber of the 20S CP and enhances its inhibitory activity (44). Our initial tests confirmed efficient modification of the *Mtb* β-subunit by SylP (SI Appendix, Fig. S1). Next, we characterized its effect on *Mtb* 20S CP peptidase activity. We reacted WT *Mtb* 20S CP with increasing [SylP] and quantified the fraction of modified β-subunits by intact protein mass spectrometry. Our peptidase assays revealed that partial SylP modification of the 20S CP similarly enhanced peptidase activity (Fig. 1E). Strikingly, 20S CP samples containing less than 22% modified β-subunits exhibited up to an 80% increased activity relative to the apo enzyme. At higher levels of modification, enzymatic activity decreased as active sites were progressively occluded by inhibitor binding (Fig. 1E). Together, these results demonstrate that substoichiometric engagement of catalytic sites, independent of inhibitor chemistry, activates the proteasome, implying positive cooperativity between neighbouring β-subunits, as observed in classical allosteric systems such as *E. coli* aspartate transcarbamoylase (46, 47).

### Investigating allostery with conventional HDX-MS

Hydrogen/deuterium exchange mass spectrometry (HDX-MS) is well suited for probing subtle changes in conformational dynamics that underlie allosteric coupling between neighbouring β-subunits within the 20S CP. The HDX rate of backbone amides reports on conformational dynamics in solution (48–50). The rate of exchange of a given amide is dependent on solvent accessibility, participation in hydrogen bonds and fluctuations in local protein structure. For example, buried or rigid regions of a protein will undergo HDX slower compared to exposed or highly flexible protein segments (48–50). HDX-MS therefore enables direct comparison of conformational dynamics between unbound and ligand-bound states of a protein (51, 52). Ligand binding frequently manifests as a decrease in deuterium uptake in areas immediately participating in the interaction, whereas allosteric changes can induce an increase or decrease in HDX across the protein structure (53, 54). We used conventional bottom-up, continuous-labeling HDX-MS for a classic two-state comparison of deuterium uptake in the apo 20S CP with that of 20S CP fully bound to ixazomib or fully reacted with SylP. We held ixazomib at saturating concentrations throughout all stages of sample preparation to maintain 90% active site occupancy during labeling. Prior to HDX measurements, we confirmed full SylP conjugation of the 20S CP by intact protein mass spectrometry under denaturing conditions. Our HDX workflow resulted in 92% sequence coverage of the β-subunit using 50 peptides, yielding a redundancy level of 2.6. Using the statistical workflow described by Weis and coworkers (55), we determined a significance threshold of 0.19 Da for differential deuterium uptake, defining the minimum change required to classify a peptide as significantly perturbed. We visualized the differential deuterium uptake of the bound states compared to apo 20S using heatmaps (SI Appendix, Fig. S2), with uptake changes binned at 0.5 Da thresholds for changes beyond ±0.19 Da. Both ixazomib binding and reaction with SylP resulted in a significant decrease in deuterium uptake across the β-subunit compared to apo 20S. As expected, the most dramatic reductions in deuterium uptake for both states occurred in the substrate binding pockets (S1 – 4) (SI Appendix, Fig. S2). Similarly, the C terminus of switch helix II also had a decrease in deuterium uptake in accordance with the ordering of residues 90-96 of switch helix II seen in both the resting and bound states of the protein (33). While SylP had a greater protective effect compared to ixazomib, the impacted regions of the protein were largely shared between the two allosteric effectors (SI Appendix, Fig. S2). Importantly, regions outside of the switch helices and catalytic site were also affected. However, discerning allosteric impacts from those of substrate binding is a challenge due to the restrictions of current HDX-MS methodology.

Bottom-up HDX-MS quantifies differences in deuterium uptake at the peptide level following acidification to quench exchange, a step that denatures the protein and dissociates multi-subunit assemblies prior to proteolytic digestion, enabling spatially resolved analysis of conformational dynamics (51, 56, 57). In symmetric homomeric assemblies, individual subunits within a single particle can occupy distinct ligation states, yet peptides derived from these sequence-identical subunits are indistinguishable by mass. While covalent modification introduces a detectable mass shift in a small number of peptides, noncovalent ligand binding and allosteric conformational changes leave no such signature. Consequently, conventional HDX-MS workflows report ensemble-averaged dynamics and cannot assign conformational responses to specific subunits in partially liganded complexes. Resolving how engagement of a subset of β-subunits is associated with conformational changes across the symmetric assembly therefore requires an approach capable of breaking β-subunit degeneracy and achieving subunit-resolved analysis.

### Stable isotope coding and assembly of hybrid complexes

To resolve the impact of effector binding in neighbouring subunits within a single 20S CP, it is imperative to discern unbound and bound β-subunits originating from a single complex. To achieve this, we assembled hybrid particles containing wild-type β-subunits (β_WT_) and a catalytically inactive variant (β_T1A_), in which the nucleophilic Thr1 was replaced with an Ala residue. Because β_T1A_ lacks the catalytic Thr1 residue, it cannot engage substrates or covalent inhibitors and therefore serves as a reporter for substrate-free subunits. Aside from this single-residue substitution at the N terminus, the two β-subunits are identical in sequence, precluding their differentiation by conventional peptide-based MS analysis. To circumvent this issue, we employed stable isotopic coding of the two subunits to distinguish them based on mass. Specifically, β_WT_ was expressed with ^15^N enrichment, while β_T1A_ was produced with natural isotopic abundance. Because ^15^N-labelled and natural-abundance peptides are chemically identical, they co-elute during LC separation and have the same drift time in the ion mobility dimension, yet are readily resolved in mass spectra. This mass difference enables direct, subunit-resolved measurement of HDX behaviour for both β_WT_ and β_T1A_ peptides within the same mass spectrum in samples where the two subunit types are intermixed (Fig. 2). Using this strategy, we incorporated isotopically coded β_WT_ and natural-abundance β_T1A_ subunits into individual 20S CPs, generating what we refer to here as “hybrid” 20S complexes for probing inter-subunit allostery.

**Figure 2.**
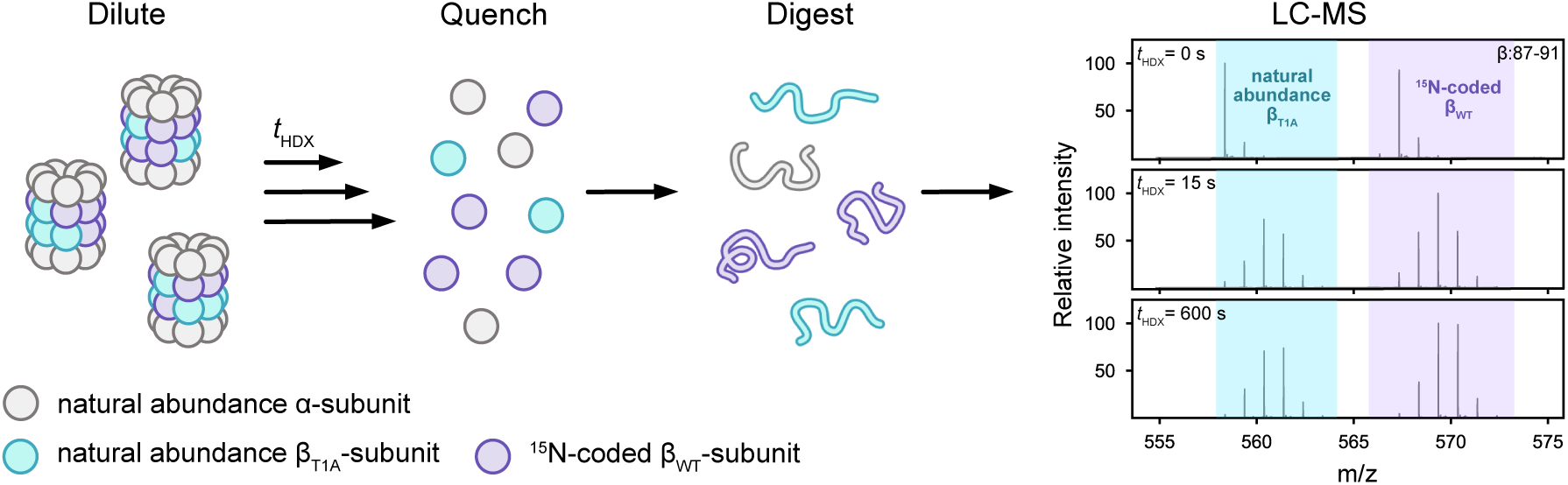
Differentially isotopically coded β-subunits form hybrid 20S CPs. ^15^N-incorporation within the β_WT_-subunit is sufficient to separate the resulting peptide spectra from those of identical sequence originating from the natural abundance β_T1A_-subunit during a typical HDX-MS workflow. MS spectra for peptide β:87-91.

To assemble these hybrid complexes, natural-abundance α- and β_T1A_-subunits and ^15^N-labelled β_WT_-subunits were individually expressed and purified, combined at defined ratios, and incubated under conditions that support 20S CP assembly (SI Appendix, Fig. S3). To assess the incorporation of β_T1A_ and β_WT_ subunits following assembly, we quantified the overall β_T1A_: β_WT_ ratios present in each sample by trypsin digestion followed by LC-MS analyses. These analyses showed that the true ratio of β_T1A_: β_WT_ in solution was consistent with those that were intended during the assembly process (SI Appendix, Fig. S4A). As an orthogonal functional validation, we measured the peptidase activity of each sample which decreased linearly with a decreasing population of β_WT_-subunits, consistent with the expected dependence of catalytic activity on the abundance of unmodified active β-subunits (SI Appendix, Fig. S4B). Together, these analyses establish the robust formation and the expected functional behaviour of isotopically coded hybrid 20S CPs, providing a validated platform for subunit-resolved HDX-MS analysis of inter-subunit allostery.

### Hybrid particle populations and probabilistic modeling of subunit composition

Although the 20S CP contains two stacked β-rings, which in principle could enable inter-ring allosteric coupling, prior studies of archaeal proteasomes have reported minimal cooperativity between opposing rings (29). We therefore build our probabilistic model as a first-pass simplification that treats each β-ring as the dominant unit of cooperativity, in which nearest-neighbour intra-ring interactions are expected to dominate. This simplification provides a tractable framework for interpreting the hybrid ensembles below, but it does not preclude the detection of cross-ring effects: if a peptide reports allosteric coupling that depends on the composition of the opposing ring, that signal will manifest as a deviation from intra-ring expectations and can be identified empirically. We return to this point later in the Results section. At any given bulk β_T1A_: β_WT_ ratio, stochastic assembly of the heptameric β-ring produces a distribution of hybrid particles that differ in subunit stoichiometry. Although the overall β_T1A_: β_WT_ ratio in each sample is experimentally defined, individual 20S CPs assemble independently and therefore populate an ensemble of distinct hybrid particles. To interpret our HDX-MS across these ensembles, we developed a probabilistic model to predict the population of hybrid particles formed at each subunit ratio. We consider a heptameric β-ring assembled from two subunit types: β_T1A_ and β_WT_. For generality, these are denoted as subunits A (β_T1A_, natural abundance, “unlabelled”) and B (β_WT_, isotopically coded, “labelled”), present at bulk fractional abundances 1 − *p* and *p*, respectively. Assuming stochastic incorporation and cyclic symmetry of the ring, each subunit position represents an independent Bernoulli trial with probability *p* of incorporating subunit B. Defining an indicator variable *X*_*i*_ ∈ {0,1} for each of the seven positions, where *X*_*i*_ = 1 denotes incorporation of subunit B, the total number of B subunits in a ring is given by:

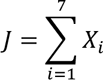

Stochastic assembly generates many positional microstates in which labelled and unlabelled subunits occupy specific sites around the ring. For example, for a ring containing a single B subunit, the configurations 0100000 and 0010000 represent distinct positional microstates in which the B subunit occupies position 2 or position 3, respectively, while the remaining positions are occupied by A subunits. The total number of such arrangements containing exactly *j* B subunits is given by the binomial coefficient 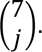 Because the heptamer is cyclic, however, configurations that differ only by rotation are physically indistinguishable. Experimentally resolvable states therefore correspond to symmetry-distinct configurations, obtained by grouping rotationally equivalent microstates. We denote the number of such rotation-unique configurations as *N*_*j*_ which is smaller than 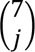 due to this symmetry reduction. This symmetry reduction is summarized in Table 1, which enumerates the number of labelled positional microstates for each value of *j*, the corresponding symmetry-equivalent configurations, and the number of rotation-unique patterns *N*_*j*_.

**Table 1.** The number of rotationally unique β-ring conformations upon β_T1A_: β_WT_ hybridization.

| $j$<br>(number of B subunits) | $\binom{7}{j}$ Labeled microstates | Symmetry-equivalent configurations | Rotation-unique patterns, $N_j$ |
| --- | --- | --- | --- |
| 0 | 1 | 1 | 1 |
| 1 | 7 | 7 | 1 |
| 2 | 21 | 7 | 3 |
| 3 | 35 | 7 | 5 |
| 4 | 35 | 7 | 5 |
| 5 | 21 | 7 | 3 |
| 6 | 7 | 7 | 1 |
| 7 | 1 | 1 | 1 |

Under stochastic assembly, the probability of forming a ring containing exactly *j* B subunits depends only on the number of such positional arrangements and their individual probabilities, yielding the binomial distribution:

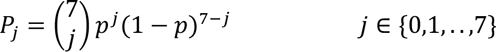

This distribution defines the relative populations of hybrid particles across the ensemble.

While *P*_*j*_ describes the global composition of β-rings in terms of A (β_T1A_) and B (β_WT_) subunits, the same framework also defines the local environments experienced by individual β-subunits. Each β-subunit has two nearest neighbours, and under stochastic assembly the identity of each neighbour (A or B) is independent and determined solely by the bulk fraction *p* of B (β_WT_) subunits. The probability that a given β-subunit has *k* B-type (β_WT_) neighbours with *k* ∈ {0,1,2} is therefore given by simple binomial statistics:

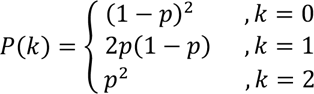

These probabilities are independent of the global ring stoichiometry and provide a direct framework for interpreting local allosteric environments experienced by individual β-subunits. Overall, this model relates bulk mixing ratios to the local environments experienced by individual β-subunits, which is critical for interpreting subunit-resolved HDX measurements.

### Subunit-resolved HDX-MS of hybrid 20S CPs

Guided by the model developed above, we assembled hybrid particles spanning a broad range of β_T1A_: β_WT_ ratios to sample the full diversity of hybrid particle stoichiometries, together with homogeneous β_T1A_ and β_WT_ 20S CP controls (Fig. 3). To further isolate specific allosteric scenarios, we included two extreme conditions with 11% and 93% β_WT_ content, where the ensemble collapses to a small number of dominant species. For example, at 93% β_WT_, ∼60% of the β-rings are fully wild-type, while ∼30% contain six β_WT_ subunits and a single β_T1A_ subunit (Fig. 3). Because the two subunit types are distinguishable by our isotopic coding strategy, the majority of β_T1A_ signal in this sample arises from isolated β_T1A_ subunits surrounded by wild-type neighbours, providing a particularly clear readout of inter-subunit allostery. Conversely, the 11% β_WT_ condition reports on the behaviour of isolated β_WT_ subunits embedded within a β_T1A_-dominated environment (Fig. 3). Together, these five mixing ratios enable systematic investigation of cooperativity across a well-defined ensemble of hybrid particle populations.

**Figure 3.**
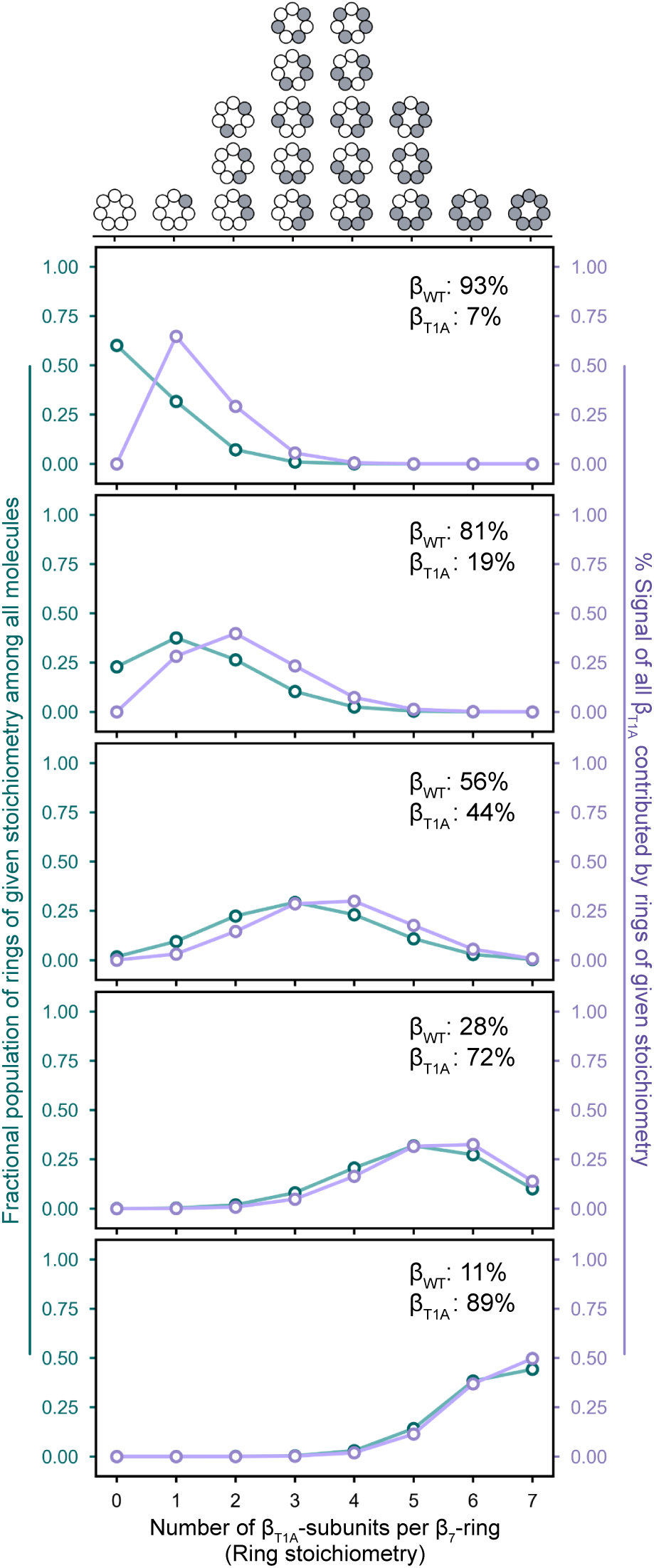
Hybridization produces a range of β-heptamer compositions, the populations of which are controlled by the ratio of β_WT_: β_T1A_. Given that the heptamer is cyclic, the number of distinguishable configurations is reduced from the total number of unique positional microstates, defined by the binomial coefficient 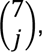 due to the inability to experimentally resolve symmetrical conformations. Using probabilistic modelling, the fractional population of each of the seven distinguishable heptamer compositions are then determined using the experimentally defined ratio of each subunit type. The signal from a given subunit type (e.g.: β_T1A_) within each of these ring stoichiometries can similarly be calculated, relating the measured uptake values of that subunit type to the local environments that it experiences.

We performed peptide mapping using an MS^E^ workflow, achieving sequence coverages of 98% for the α-subunit, 91% for β_T1A_, and 90% for β_WT_ with redundancy levels of 3.6, 3.3, and 3.1, respectively. As above, we used the statistical analysis described by Weis (55), which yielded a significance threshold of 0.37 Da, above which differences in deuterium uptake were considered significant. We visualized differential deuterium uptake using heatmaps. Building on our previous HDX-MS work demonstrating that orthosteric inhibition or mutation of βThr1 alters 20S CP dynamics (33), we used the reaction of isotope-coded hybrid particles with SylP to simultaneously measure local effects within ligand-bound β_WT_ subunits and the allosteric responses occurring in neighbouring β_T1A_ subunits. Allosteric effects within β_WT_-subunits are present but are partly masked by the strong local protection induced by SylP engagement; the β_T1A_ subunits, which cannot bind SylP, therefore provide a cleaner readout of pure allosteric coupling. We first quantified the direct effect of allosteric effector engagement by comparing apo and SylP-reacted states of β_WT_ subunits at matched β_WT_ fractions (Fig. 4, SI Appendix, Fig. S5). Across all ratios, SylP reaction caused a near-global reduction in conformational dynamics, similar to our earlier observations in homogeneous 20S CP (SI Appendix, Fig. S2), with the substrate-binding pockets and switch helices showing the strongest protection. Increasing incorporation of β_T1A_ subunits attenuated protection in some regions (e.g., β87-98) while enhancing it in others (e.g., β214-238) (Fig. 4). The intermediate hybrid particles with 28%, 56%, and 81% β_WT_ content showed the largest number of peptides with significant differential uptake, indicating maximal divergence between apo and holo conformational dynamics at these stoichiometries.

**Figure 4.**
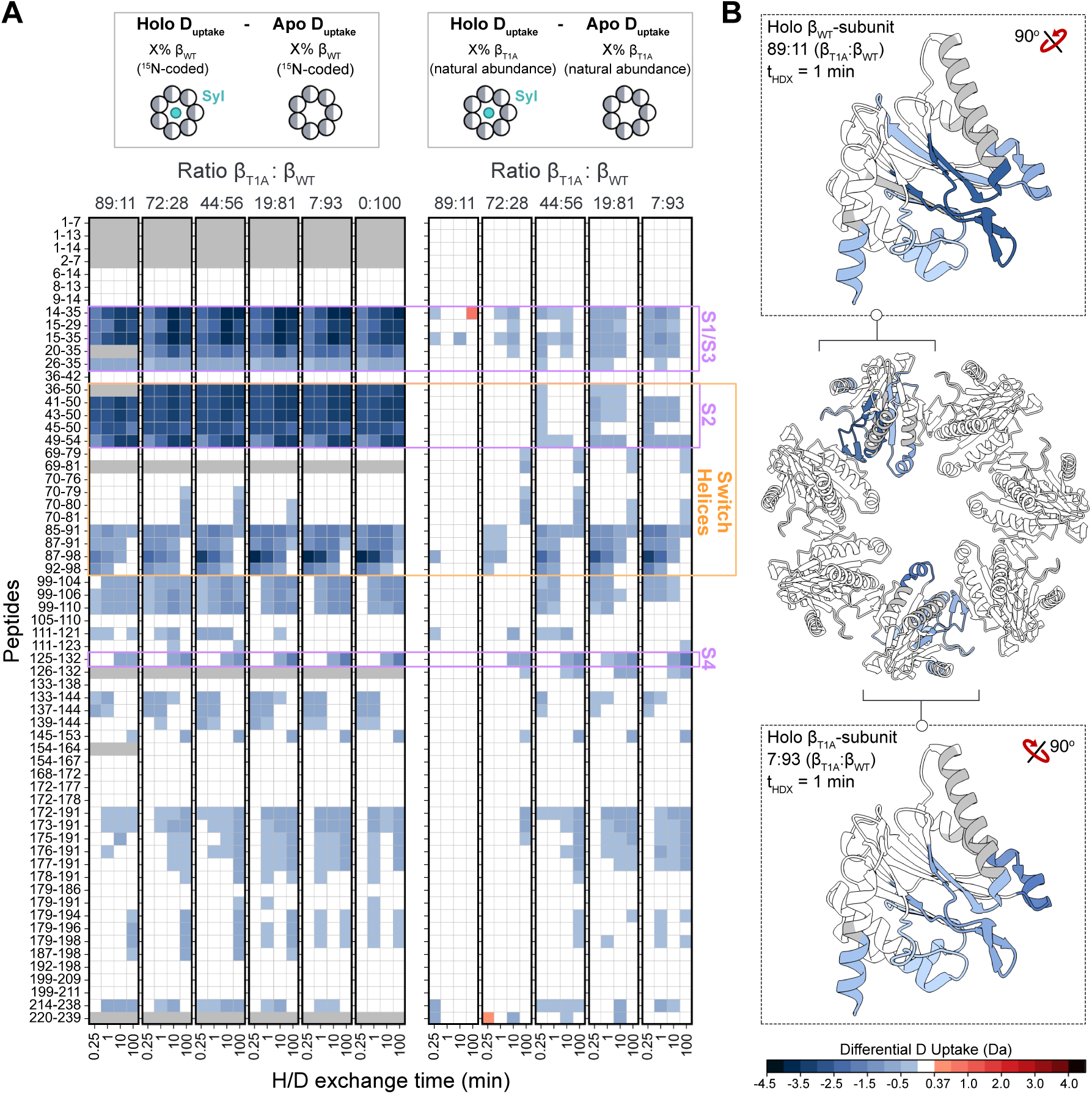
Ligand binding allosterically impacts neighbouring, unbound protomers. (**A**) Heatmaps depicting the relative deuterium uptake of each SylP-reacted hybrid complexes compared to their unbound equivalent for both the β_WT_- (left) and β_T1A_-subunits (right). Gray boxes indicate absent data. Peptides associated with the active site (S pockets) and switch helices are indicated; and (**B**) The relative deuterium uptake at *t*_HDX_ = 1 min projected onto the structure for the 89:11 (β_T1A_: β_WT_) ratio of the β_WT_-subunit (top) and the 7:93 ratio of the β_T1A_-subunit (bottom). Statistical analysis determined a significance interval of 0.19 Da which was used as the lower bound. All other changes were determined to be significant above 0.5 Da, with differences colour-coded onto the structures. Uncovered regions are coloured gray. All structures are visualized in Chimera 1.7 (PDB 9ce5).

We next tested whether ligand binding to β_WT_ subunits induced allosteric responses in neighbouring β_T1A_ subunits. Crucially, this analysis is only possible because our isotopic coding strategy enables direct, subunit-resolved measurement of HDX behaviour within a single hybrid complex. Despite their inability to bind ligand directly, β_T1A_ subunits in every hybrid population showed decreased deuterium uptake upon SylP reaction of the complex (Fig. 4; SI Appendix, Fig. S5). As observed for β_WT_, the most strongly protected regions included the substrate-binding pockets and switch helices, with higher β_WT_ fractions generally producing greater protection (e.g., β:87–98). In a given hybrid population, up to 35 β_T1A_ peptides displayed altered uptake, and the same structural regions emerged across both β_WT_ and β_T1A_ subunits (Fig. 4). These included sites near the catalytic residues and switch helices, as well as more distal segments spanning residues 172-191 which encompasses a loop in proximity to the C-terminal α-helix and neighbouring β-subunit (Fig. 4). Thus, ligand binding to one β-subunit is sufficient to remodel the conformational dynamics of adjacent, binding-incompetent subunits, identifying the specific structural elements that mediate cooperative allosteric coupling across the proteasome β-ring.

In our previous study, we demonstrated that cooperativity within the *Mtb* 20S CP is sensitive to substitutions within the α- or β-subunits (33). In particular, the β_T1A_ single amino acid substitution proved to stabilize the inactive state of the 20S CP which we characterized using a combination of HDX-MS and electron cryomicroscopy (33). Building on this foundation, our isotope-resolved hybrid system enabled us to determine how co-assembly of β_WT_ and β_T1A_ subunits reshapes the conformational landscape of each subunit within the same particle. To do so, we compared apo hybrid samples to their respective homogeneous controls (20S_WT_ and 20S_T1A_). In β_WT_ subunits, hybridization increased conformational plasticity, with the number of affected peptides rising as the fraction of β_T1A_ increased (Fig. 5; SI Appendix, Fig. S6). At β_WT_ fractions between 11% and 81%, peptides within the substrate-binding pockets showed increased deuterium uptake, while changes in the switch helices peaked at 28% and 56% β_WT_. Peptides spanning residues 171-194, corresponding to a loop near both the C-terminal α-helix and neighbouring β-subunit, showed reduced protection at 11-56% β_WT_, with the largest changes occurring at the lowest β_WT_ fraction (Fig. 5A). Across all hybrid samples, including the 93% β_WT_ condition, which otherwise showed minimal perturbation, the C-terminal peptide β214-238 which forms the C-terminal α-helix consistently exhibited altered dynamics. Hybridization exerted an even stronger effect on β_T1A_ subunits, with up to 23 peptides displaying altered uptake in a given sample (Fig. 5). All hybrid populations showed increased dynamics in the substrate-binding pockets and switch helices relative to 20S_T1A_, while higher β_WT_ fractions induced decreased dynamics in additional regions (e.g., β133-144), with the strongest protection observed at the highest β_WT_ content (Fig. 5A). Strikingly, the 56% β_WT_ condition, where the two subunits are most evenly distributed, produced the largest number of affected peptides, revealing maximal sensitivity to inter-subunit allosteric coupling at intermediate stoichiometries. Similarly, reaction with SylP and the hybridization of β_WT_-and β_T1A_-subunits also modulated the conformational plasticity of the 20S CP. These changes are allosterically transmitted to the α-subunit in accordance with our previous findings (SI Appendix, Fig. S7).

**Figure 5.**
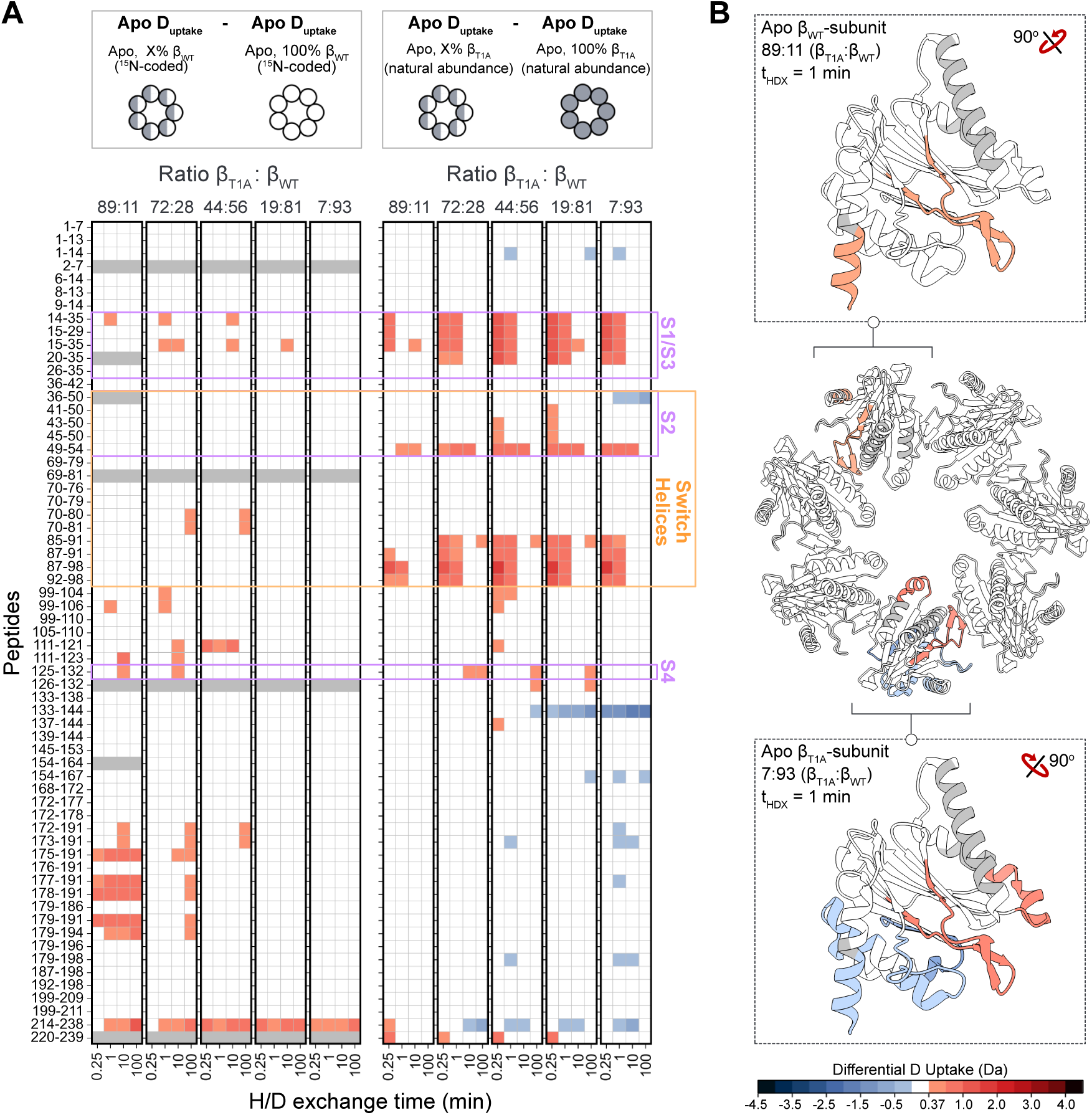
Hybridization impacts the conformational dynamics of both β-subunit types. (**A**) Heatmaps for each tested β_WT_: β_T1A_ ratio depicting the relative deuterium uptake of the apo, hybridized β_WT_-subunit compared to the 20S_WT_ control (left) and for the apo, hybridized β_T1A_-subunit compared to the 20S_T1A_ control (right). Gray boxes indicate absent data. Peptides associated with the active site (S pockets) and switch helices are indicated; and (**B**) The relative deuterium uptake at *t*_HDX_ = 1 min projected onto the structure for the 89:11 (β_T1A_: β_WT_) ratio of the β_WT_-subunit (top) and the 7:93 ratio of the β_T1A_-subunit (bottom). Statistical analysis determined a significance interval of 0.19 Da which was used as a lower bound. All other changes were determined to be significant above 0.5 Da, with differences being coloured according to the colour bar. Uncovered regions are coloured gray. All structures are visualized in Chimera 1.7 (PDB 9ce5).

### Hybridization identifies inter-β allosteric pathways within the 20S CP

The hybrid particle sample series provides a second analytical axis beyond H/D exchange time. At any chosen H/D exchange time, the relative deuterium uptake of a given peptide can be plotted as a function of the β_WT_ content of the hybrid ensemble, and the slope of this dependence reports on whether the peptide’s conformational dynamics are coupled to ring composition. We refer to this slope, fitted independently at each H/D exchange time, as the compositional slope. Peptides whose conformational dynamics are independent of composition have compositional slopes close to zero; peptides that participate in inter-subunit cooperative coupling have systematically non-zero slopes. Because the isotopic coding strategy resolves the same peptide separately through the β_T1A_ and β_WT_ subunits of the hybrid particle, we obtain two independent compositional slopes per peptide and plot these against one another (Fig. 6, SI Appendix, Fig. S8). A peptide whose compositional slope crosses the global 95 % confidence interval in the β_T1A_ subunits only is classified as “β_T1A_-significant”; in the β_WT_ subunit only, as “β_WT_-significant”; and in both subunits, as “co-significant”. We applied this framework to two complementary comparisons. The first contrasts the SylP-reacted and apo states of each hybrid complex (Fig. 6A; SI Appendix, Fig. S8A). Because every β_WT_ subunit in a SylP-reacted sample is itself bound, a non-zero compositional slope in this comparison reports on how ligand engagement propagates from bound β_WT_ subunits to their neighbours. The second comparison contrasts each apo hybrid with its homogeneous apo control, 20S_WT_ or 20S_T1A_ (Fig. 6B; SI Appendix, Fig. S8B). No ligand is present on either side of this comparison, so a non-zero compositional slope reports purely on dynamics that are sensitive to neighbour identity in the absence of ligand. Together, the two comparisons partition the inter-subunit signal into binding-driven and composition-driven contributions.

**Figure 6.**
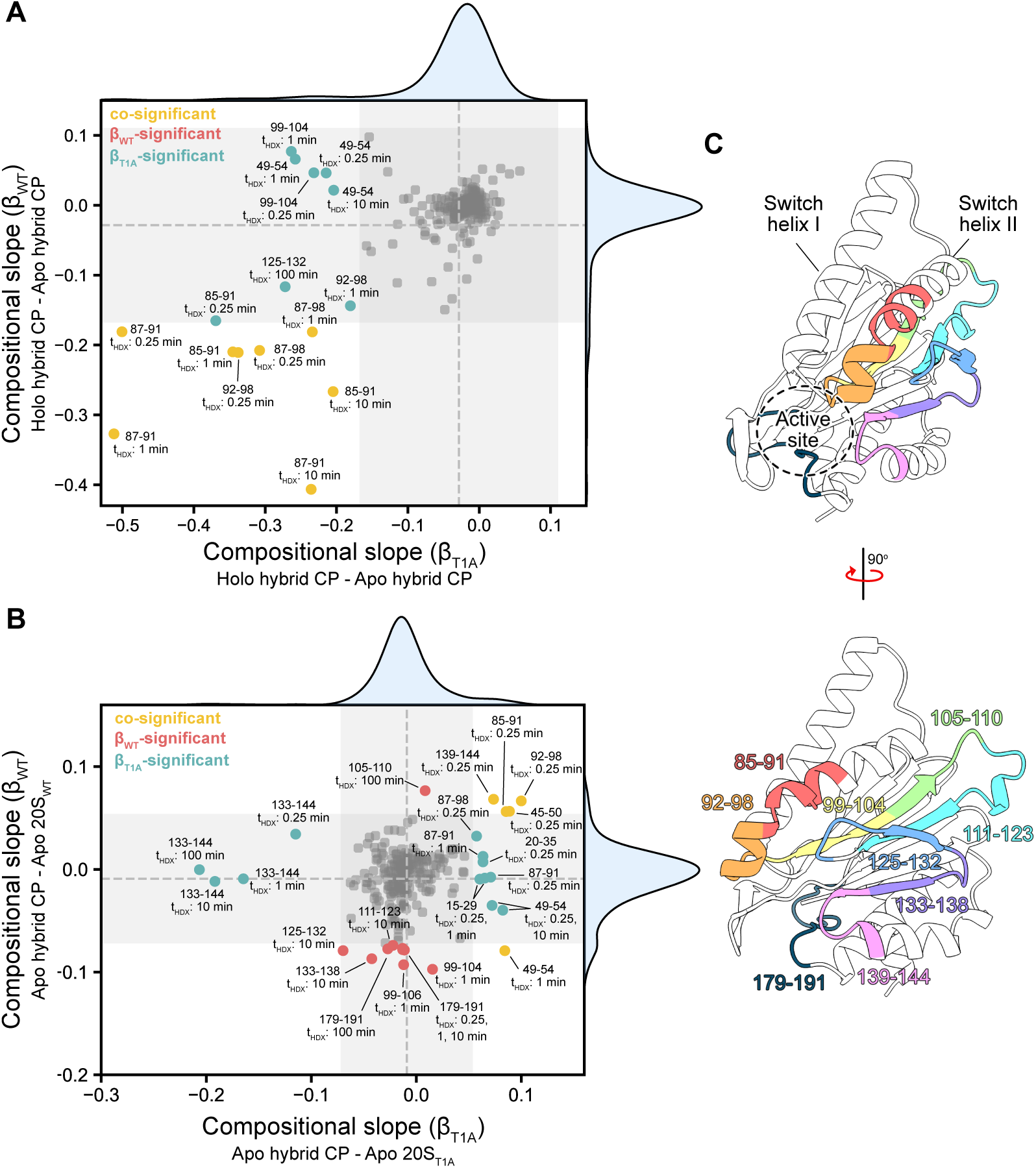
Measuring deuterium uptake across varying hybrid compositions identifies regions of allosteric communication. (**A**) Volcano plot in which each point represents the compositional slope of the change in relative deuterium uptake of a peptide at a given H/D exchange time in response to the titration of the SylP-reacted β_WT_-subunit; (**B**) Volcano plot in which each point represents the compositional slope of the change in relative deuterium uptake in response to the titration of the unbound β_WT_-subunit, compared to the 20S_WT_ or 20S_T1A_ controls for the β_WT_- and β_T1A_-subunits, respectively. Peptides are coloured based on the datasets in which they are deemed significant: peptides appearing only in the β_T1A_-dataset are shown in teal, peptides appearing only in the β_WT_-dataset are shown in red, and peptides appearing in both are shown in yellow. The change in relative uptake for each peptide is compared to the globally calculated mean (see Suppl. Fig 8) of the respective dataset. The calculated mean is shown as gray dashes and the 95% C.I. is shown as gray boxes across both axes, respectively. Histograms show the population density of peptides for the β_T1A_- (top) and β_WT_-subunits (right); and (**C**) The β-subunit with the switch helices, active site, and peptides identified in this analysis highlighted. Structure visualized in Chimera 1.7 (PDB 9ce5).

In the first comparison, the majority of peptides showed compositional slopes close to zero, as expected. The clearest exception was the C terminus of switch helix II (β:85-91, 87-91, 87-98, 92-98), which displayed strong negative slopes in both subunit types and was therefore co-significant, consistent with progressive rigidification of this region as more β_WT_ subunits become SylP-bound. The co-significance of switch helix II is mechanistically informative. Because every β_WT_ subunit in a SylP-reacted hybrid is itself bound, any β_WT_ signal arising purely from direct ligand engagement should saturate at the holo-apo difference of an isolated β_WT_ subunit and not vary with composition. A non-zero compositional slope in the β_WT_-derived peptides of switch helix II therefore indicates that even a fully bound β_WT_ subunit additionally responds to the binding state of its β_WT_ neighbours within the same particle, a direct signature of inter-β_WT_ cooperativity in the saturated state. Two β_T1A_-significant peptides spanning the S2 and S4 pockets (β:49-54 and 125-132) also carried slopes in this comparison; in their case, β_T1A_ cannot bind SylP, so the slope is unambiguously allosteric. No β_WT_-significant peptides were detected in these regions, again consistent with the saturation argument above. The β_T1A_ subunits therefore serve as the primary, but not exclusive, reporter for inter-subunit allosteric transmission in this comparison: they isolate allosteric signals from local binding effects, while the co-significant peptides provide an independent confirmation that the same allosteric pathway remains observable within the saturated β_WT_ subunit.

We next applied the analysis to the apo-vs-apo composition comparison. Switch helix II again emerged in both subunit types (β:85-91, 92-98), as did peptides spanning the S2 pocket (β:45-50, 49-54), confirming that even unbound β_WT_ subunits sense the identity of their neighbours. This comparison also revealed peptides outside the previously identified allosteric hotspots. In the β_WT_ subunit, a contiguous set of peptides spanning the four β-strands immediately preceding switch helix II showed significant negative slopes (β:99-104, 99-106, 105-110, 111-123, 125-132, 133-138, 139-144; Fig. 6B,C). Spatially, these peptides occupy the interface between switch helix II of one subunit and the active site of its intra-ring neighbour. To rationalize how this segment could mediate inter-β communication, we inspected the corresponding β:β interface in the structure and identified three direct inter-subunit contacts within the β7-ring: G128:A50′, E133:D30′, and L144:M25′. Each of these residues is also positioned within *ca.* 4 Å of an S-pocket residue on the partner subunit. G128 contacts the S4 residue D124; M25 contacts T21 and Q22 of the S3 pocket; and D30 contacts R18, the residue immediately preceding the catalytic D17. These contacts therefore constitute a plausible structural conduit linking switch helix II of one β-subunit to the active site of its lateral neighbour.

We identified a second, distinct allosteric pathway through β:179-191. This peptide was the most consistently composition-dependent in the β_WT_ subunit, with a significant negative slope at every HDX timepoint (Fig. 6B,C). Unlike the intra-ring peptides above, β:179-191 sits at the interface between opposing β-rings, and its composition-dependent behaviour therefore refines the single-ring framework introduced earlier: in addition to the dominant intra-ring pathway, the hybridization HDX-MS approach is sufficiently sensitive to detect inter-ring β: β coupling at a defined structural location. Within its own subunit, β:179-191 contacts residues within or near the S pockets (D17, R18, R19, R29). Across the inter-ring interface, R188, V187, and S179 form direct contacts with D176, S222, and N24/I26 of the partner-ring subunit, several of which lie in the vicinity of that subunit’s S3 pocket. Inter-β allosteric coupling within the *Mtb* 20S CP therefore appears to operate along two structurally distinct routes: a lateral pathway connecting switch helix II to the active site of the adjacent intra-ring subunit, and an axial pathway connecting β:179-191 to the S pockets of the opposing ring.

Finally, we examined the β_T1A_ subunit under the same composition comparison. Beyond the switch helix II and S2-pocket peptides that co-significantly tracked composition in both the β_WT_ and β_T1A_ subunits, β_T1A_ also showed significant positive slopes in S1 and S3 peptides containing the active-site residues D17 and K33 (β:15-29, 20-35). These regions become more flexible as β_T1A_ subunits are replaced by β_WT_ neighbours. We interpret this as a partial rescue of active-site dynamics: the β_T1A_ substitution stabilizes the inactive conformation in homogeneous 20S_T1A_ (33), and incorporation of β_WT_ neighbours restores more wild-type-like flexibility. Outside the active site, the only β_T1A_-significant peptides mapped to β:133-144 and β:139-144, the same intra-ring interface segment implicated in the β_WT_ subunit. The peptide β:133-144 in particular showed significant slopes at every timepoint tested, identifying this segment as the dominant intra-ring conduit for composition-driven changes in β_T1A_. Notably, this peptide also contains S141, recently proposed to participate in a catalytic pentad with D17, K33, and D178 (58) (see the Discussion).

## Discussion

Cooperativity within the degradation chamber of proteasome systems is incompletely understood. Early characterization of the eukaryotic proteasome system using chemical inhibitors found that the catalytic subunits were allosterically linked (35–37), however, whether such relationships were present in the bacterial homologue remained unknown. Here, we show that such relationships also exist between catalytic sites of the prokaryotic *Mtb* 20S CP. We discovered that engagement of orthosteric inhibitors to a subset of β-subunits paradoxically stimulated enzymatic activity. We interpreted these data in the context of allostery where effector binding promotes the active conformation in both the effector-bound and -unbound subunits via inter-subunit cooperativity. This mechanism has previously been observed in other proteases (59–61). However, investigating these allosteric relationships is complicated by the symmetric and homomeric β-chamber of the *Mtb* proteasome system. The issue of subunit degeneracy has previously been overcome in nuclear magnetic resonance (NMR) spectroscopy studies through differential isotopic labelling to isolate signals from a given subunit type against a varied perdeuterated NMR-invisible background (41, 62–64). We were also inspired by pioneering HDX-MS studies that exploit isotopic enrichment to separate the signal spectra of asymmetric protein complexes composed of isoforms (65) or proteins with high sequence homology (66). Incorporating these methodologies, we isotopically coded β-subunits to probe the allosteric impacts of ligand binding in hybridized *Mtb* 20S CPs.

Binding of a substrate-mimic induced extensive changes within the conformational dynamics of the β_WT_-subunit in both pure and hybridized particles. These same regions were similarly impacted in hybridized β_T1A_-subunits through allosteric modulation, with differential uptake frequently correlating to increasing presence of reacted β_WT_-subunits. Protection within the S pockets and switch helices substantiate that the active sites are allosterically linked and further highlight the required disorder to order transition of C-terminal residues in switch helix II to accommodate the active conformation. Notably, while β_WT_-subunits stabilized switch helix II in β_T1A_-subunits (Fig. 4A), β_T1A_-subunits lessened protection in WT subunits suggesting that the subunits mutually impact the dynamics of the other. Both WT and variant proteins also had decreased uptake in peptides containing residues Ser141 and Asp178 (Fig. 4A). These residues were recently proposed to support the Thr1 amide for catalytic reaction and participate in a catalytic pentad alongside Asp17 and Lys33 (8, 58). Given the protection of these regions in response to inhibitor binding, our results support the structural involvement of these residues in substrate binding.

When comparing the unbound hybrid samples to either a 20S_WT_ or 20S_T1A_ control, hybridization clearly impacted the conformational dynamics of both WT and variant subunits. The S pockets had increased uptake in both proteins, with more pronounced changes in the β_T1A_-subunit. This suggests that hybridization mutually increases plasticity within the active sites of either protein, with 20S_T1A_ being more sensitive to ring composition. Interestingly, the switch helices in β_T1A_-subunits also had increased plasticity. We have previously shown that the C-terminal region of switch helix II (β:87-98) had decreased dynamics in the 20S_T1A_ compared to 20S_WT_ (33). Current results suggest that interaction with WT subunits restore more typical dynamics to this region in hybridized β_T1A_-subunits. Regions of the β_T1A_-subunits also undergo rigidification at higher β_WT_ populations. These include peptides within the protein core as well as α-helices and loops in contact with the neighbouring β-ring. Together, these results suggest that dynamics of the β_T1A_-subunits are increased between intra-ring contacts but are more anchored across inter-ring contacts. Comparatively, the β_WT_-subunit has increased plasticity between inter-ring contacts, namely the loop formed by residues 179-191 and the C-terminal helix of the opposing subunit in which it packs (Fig. 5B). This empirically validates the modest β:β inter-ring coupling that our single-ring framework was constructed to remain agnostic about. Interestingly, the largest number of differentially impacted regions was not exclusively correlated with the extreme β_T1A_: β_WT_ ratios. In both bound and apo samples, the intermediate ratios had additional changes in dynamics to regions both within and in contact with the N terminus of switch helix II.

In the current study, we presented isotopic enrichment in HDX-MS as a powerful strategy for dissecting symmetry-related allosteric mechanisms in large multi-subunit assemblies. Differential ^15^N-labelling separated peptide signals from natural-abundance counterparts of identical sequence, enabling subunit-resolved HDX-MS measurements within a single hybrid complex. Our methodology generalizes to any homo-oligomeric assembly whose subunits can be co-mixed in an isotopically resolvable manner, and the approach turns the symmetry that ordinarily masks allostery into a tunable analytical variable. Several constraints define the systems to which the approach is most readily applied. First, the method requires an amino acid substitution that disables effector binding in one subunit type, such as the T1A substitution we exploited here. Such mutations are not known for every system, and when known they may still fail to co-assemble with wild-type subunits into a single complex. Second, controlled co-assembly is itself non-trivial. Some systems require denaturing and refolding to drive subunit incorporation, others do not tolerate this treatment, which may lead to insoluble protein. Third, stochastic mixing always produces a distribution of subunit stoichiometries at any given bulk ratio rather than a single, well-defined composition. The clearest exception arises at highly skewed mixing ratios, where nearly pure single-incorporated or non-incorporated particles dominate the ensemble, but those compositions introduce their own measurement challenges. The minor subunit is then present at very low abundance against a large excess of the other, straining the dynamic range of the MS readout. Extending the approach to systems that lack a binding-incompetent variant, to assemblies that resist controlled co-mixing, and to compositions at the extremes of the mixing range will require complementary strategies such as designed weak-binding analogues and isolation of a distinct subunit under HDX quench conditions. With these refinements, hybridization HDX-MS provides both a structural basis for investigating homologous complexes and mechanistic insight into the broader class of symmetric multi-subunit assemblies whose function depends on inter-subunit communication.

## Materials and Methods

Details of gene expression, protein purification, and biochemical measurements, along with mass spectrometry experiments are provided in the SI Appendix.

## Data, Materials, and Software Availability

Mass spectrometry data are available from the MassIVE database as entry MSV000102181. All other data are included in the manuscript and/or supporting information.

## Acknowledgments

M.T. acknowledges support from a Natural Sciences and Engineering Research Council of Canada Postgraduate Doctoral Scholarship. Financial support was provided by a Canadian Institutes of Health Research Project Grant PJT451412 to S.V., and by the Natural Sciences and Engineering Research Council of Canada grants RGPIN-2021-02843 to S.V and CRD-PJ 480432 to D.J.W. T.B. acknowledges funding from the Austrian Science Fund (FWF) Cluster of Excellence COE7 ‘Microbiomes drive Planetary Health’ [doi.org/10.55776/COE7].

