## Supplementary material for "Resolving symmetry-masked allosteric cooperativity in the *M. tuberculosis* proteasome core particle": SI Appendix

**Affiliations:**
[1]Department of Molecular and Cellular Biology, University of Guelph, Guelph, Ontario N1G 2W1 Canada

[2]Faculty of Chemistry, Institute for Biological Chemistry & Centre for Microbiology and Environmental Systems Science, Department of Microbiology and Ecosystems Science, Josef-Holaubek-Platz 2 (UZA II), University of Vienna, 1090 Vienna, Austria

[3]Department of Chemistry, York University, Toronto, Ontario M3J 1P3, Canada

[4]Waters Corporation, Stamford Avenue, Wilmslow SK9 4AX, UK

**This PDF file includes:**

Supporting text
Figures S1 to S9
Tables S1 and S2
SI Appendix References

### Materials and methods

#### Plasmids, constructs, and protein purification

Protein expression and purification was carried out as described in (1). Briefly, the genes encoding the α- (Uniprot: P9WHU1, prcA, Rv2109c) and β-subunit lacking the propeptide (Uniprot: P9WHT9, prcB, Rv2110c) of *Mycobacterium tuberculosis* proteasome core particle (CP) were synthesized and inserted into separate pET24a vectors encoding either an N-terminal or C-terminal cleavable His-SUMO tag, respectively. A T1A point mutation was introduced to the β-subunit using the Quikchange method.

Plasmids were individually transformed using the heat-shock method into chemically competent T7 Express *Escherichia coli* cells or BL21(DE3) cells to produce natural-abundance or isotopically labelled protein, respectively. T7 Express cells were then grown in lysogeny broth (LB) media, while minimal media (M9) supplemented with $^{15}$N-labelled ammonium chloride (Cambridge isotope laboratories, inc., MA, USA) was used to produce isotopically labelled protein. All media was supplemented with 30 µg/mL kanamycin. Cultures were grown at 37 °C prior to induction with 1 mM isopropyl β-D-1-thiogalactopyranoside (IPTG) at an $OD_{600}$ of 0.6. Protein expression proceeded for 18 h at 16 °C before harvesting cells by centrifugation at 4,000 ×*g* for 15 min at 4 °C. The cells were then resuspended in lysis buffer (300 mM NaCl, 20 mM imidazole and 50 mM Tris-HCl, pH 7.0) and lysed using an Emulsiflex-C3 high-pressure homogenizer (Avestin Inc., Ottawa, Canada). The resulting homogenate was clarified by centrifugation at 23,700 ×*g* for 50 min at 4 °C, of which the supernatant was passed over a $Ni^{2+}$-charged chelating FastFlow™ Sepharose column. The column was subsequently washed with 40 mL of lysis buffer, and 30 mL of lysis buffer supplemented with 50 mM imidazole before finally eluting the protein using lysis buffer supplemented with 500 mM imidazole.

To produce hybrid CPs, the concentration of elution fractions of isotopically-labelled $\beta_{WT}$- and natural abundance $\alpha_{WT}$- and $\beta_{T1A}$-subunits were determined spectrophotometrically (guanidine hydrochloride-denatured protein). Extinction coefficients of 17,880 $M^{-1}cm^{-1}$ for the His-SUMO-$\alpha_{WT}$-subunit, and 21890 $M^{-1}cm^{-1}$ for the β-subunits with C-terminal His-SUMO tags, obtained from Expasy ProtParam web-based tool (https://web.expasy.org/protparam/), were used to calculate concentrations. The $\beta_{WT}$- and $\beta_{T1A}$-subunits were then combined at defined ratios prior to mixing with the α-subunit. The different reactions were then separately dialyzed against 300 mM NaCl, 1 mM DTT, 50 mM Tris-HCl, pH 7.0 for 18 h at 4 °C in the presence of TEV and Ulp1 proteases to remove the affinity tags and to promote the assembly of the hybrid 20S CP complexes. For all complexes, the affinity tags and other impurities were then removed by passing the mixtures over the IMAC column for a second time. The resulting flow-through was concentrated using a 10 kDa MWCO Amicon Ultra-15 centrifugal filter (Millipore). The fully formed 20S CP was then separated from unassembled subunits via size-exclusion chromatography (SEC) using 100 mM NaCl and 50 mM $NaH_2PO_4$, pH 7.4 as the running buffer. SDS-PAGE was used at each step to assess protein expression and purity. The final 20S CP concentration was determined spectrophotometrically (guanidinium chloride-denatured protein) using extinction coefficients of 16,390 for the $\alpha_{WT}$-subunit, and 20,400 $M^{-1}cm^{-1}$ for the β-subunits.

#### Conjugation with SylP

To isolate the substrate-bound state, the 20S CP was reacted with the covalent inhibitor, Syringolin probe (SylP). Purified protein samples were incubated with 0 – 2-fold SylP to get partially-reacted 20S CP for functional characterization (see below). Samples were reacted with greater than 3-fold SylP to yield fully reacted core particles for both functional and structural characterization. All reactions were allowed to proceed for 18 h at 4 °C in 100 mM NaCl and 50 mM $NaH_2PO_4$, pH 7.4

with DMSO kept under 1% (v/v). Unreacted SylP was subsequently removed via buffer exchange using a 10 kDa MWCO Amicon Ultra-0.5 centrifugal filter (Millipore). Samples were washed a minimum of three times with reaction buffer at RT before recovering. The percent of modified β-subunits was then determined using intact LC-MS measurements.

1.5 - 3 pmol of each respective SylP-reacted 20S CP sample was injected onto an ACQUITY™ BEH™ C4 Column (1.7um, 2.1 x 150mm) (Waters) connected to a Waters ACQUITY UPLC™ I-Class System. Reverse-phase liquid chromatography was used to separate proteins over a 13-min $H_2O$: acetonitrile (ACN) gradient, where both mobile phases were acidified using 0.1% (v/v) formic acid (FA). Over the first 7 min, ACN was linearly increased from 2–85% before a single sawtooth gradient where it was ramped from 2–85 % over 3 minutes. A final flush of 2 % ACN was used to equilibrate for the next injection. Samples were separated using a flow rate of 200 $\mu L\ min^{-1}$ and a column temperature of 40 °C.

Eluent from the UPLC was directed to a SYNAPT™ G2-Si quadrupole time-of-flight (Q-TOF) Mass Spectrometer. The system was fitted with a standard dual emitter electrospray ionization (ESI) source which was operated in positive-ion mode with a capillary voltage of +3 kV. The TOF mass analyzer was set to resolution mode, and surveyed precursor ions from 50–2,000 Th using 0.4 second scans. The system was both externally calibrated using sodium iodide (700008892) over the 50 – 2000 m/z range and dynamically calibrated by infusing 2 µM Leucine Enkephalin (peptide sequence YGGFL; 1+ *m/z* 556.2771) dissolved in 50% (v/v) ACN and 0.1% (v/v) FA at a flowrate of 10 $\mu L\ min^{-1}$ from the LockSpray capillary, which was sampled every 20 sec.

Data analysis was carried out using MassLynx™ Software (Waters). Spectra were summed over a 0.7 min window of the chromatogram, which corresponds to the elution profile for the β-subunit. The MaxEnt1 function was then used to deconvolute using an output range of 24-29 kDa with a resolution of 1 Da/channel. A maximum number of 25 iterations were set prior to converging.

#### Functional characterization

The inhibitory effect of covalent small molecules on the peptidase function of the 20S CP were determined against the tripeptide benzyloxycarbonyl-Val-Leu-Arg-7-amino-4-methylcoumarin (Z-VLR-AMC, Genscript). Characterization of the reversible inhibitor, ixazomib (Selleck Chemicals LLC), required that both substrate and protein stocks contained 0 – 10 µM inhibitor and 1% DMSO (v/v) to avoid dilution. Substrate stocks were prepared in water while enzyme stocks were prepared in 5x reaction buffer. Reactions were then initiated via the addition of the protein/ixazomib stock. Comparatively, with SylP being an irreversible inhibitor 20S CP conjugation was completed as described above. Substrate stocks were prepared in water with 10% DMSO and enzyme stocks were prepared in 5x reaction buffer. Reactions were initiated with the addition of conjugated 20S CP.

The peptidase activity of the hybrid 20S CPs were similarly tested against Z-VLR-AMC. Substrate stocks were prepared in water with 10% DMSO (v/v) at 10-fold the intended final concentration. Hybrid particle stocks, ranging from 0 – 100% $\beta_{WT}$, were prepared in 5x reaction buffer. Reactions were initiated via the addition of the protein stock.

All reactions were performed at 25 °C in 20 mM NaCl, 10 mM $NaH_2PO_4$, pH 7.4 and 1% DMSO (v/v) with a final protein concentration of 21.4 nM and Z-VLR-AMC concentration of 150 µM. Cleavage of Z-VLR-AMC was monitored using a Cary Eclipse fluorometer (Agilent Technologies, Mississauga, Canada) at 380 nm excitation and 450 nm emission wavelengths, respectively, with a bandpass of 5 nm. A standard AMC curve was used to convert relative fluorescence values into the concentration of product formed. Initial velocities were extracted from the first 30 seconds of each reaction using Python scripts written in-house. All functional assays were carried out in technical triplicates. Data were visualized using in-house scripts written in Python v3.8.

#### Trypsin digest coupled to mass spectrometry

For combinatorically combined (hybridized) particles, the true ratio of $^{15}$N-labelled $\beta_{WT}$: natural-abundance $\beta_{T1A}$ in the sample was determined using a tryptic digest. Each protein sample was diluted to 5 µM in 5 mM DTT, 50 mM Tris-HCl, pH 8.0. All protein samples were denatured at 70 °C for 15 minutes prior to the addition of MS-grade trypsin/Lys-C (Promega) at a ratio of 1:50 (w:w) protease:20S CP. Digestion was allowed to proceed for 18 h at 37 °C before quenching with 0.1% (v/v) formic acid. Samples were stored at -80 °C prior to MS analysis.

A Waters ACQUITY UPLC I-class System fitted with an ACQUITY BEH C18 Column (1.7 µm particle size; 1 mm × 100 mm) (Waters) was used to separate 2.5 pmol of the tryptic digests by reverse-phase liquid chromatography. A 30-min $H_2O$: ACN gradient in which ACN content was increased linearly from 2 – 60% over the first 22 min then ramped to 80% in the next 3 min was used to separate peptides. Both mobile phases were acidified with 0.1% (v/v) FA. The column temperature was held at 40 °C, and the flow rate was 150 µL min$^{-1}$. Between sample injections, cleaning runs consisting of a 30-min sawtooth gradient where ACN was increased from 2 – 85% four times were used to minimize sample carryover. The UPLC outflow was directed to a SYNAPT G2-Si Q-TOF Mass Spectrometer as described above. The system was externally and dynamically calibrated as described above. The TOF mass analyzer was set to resolution mode. The sampling cone was set to 40 V, and the source offset was set to 30 V. The source block and desolvation temperatures were 80 and 600 °C, respectively.

Spectra were recorded between 50 and 2,000 m/z with a scan time of 0.4 s for data-independent acquisition. The trap TWIG collision energy was ramped between 20 – 40 V in alternating scans to induce CID. All other TWIGs were controlled automatically. Data were searched against the β-subunit sequence (UniProt: P9WHT9) using ProteinLynx Global Server v.3.5.2 (Waters). Trypsin was set as the digestion agent with an allowance of one missed cleavage. No modifications were permitted. This analysis yielded a total of 60 peptides, which were then manually inspected in DynamX v3.0 (Waters). Intensities of the surviving peptides, of which there were corresponding $^{15}$N-labelled and natural abundance sequences, were then used to determine the ratio of $\beta_{WT}$: $\beta_{T1A}$.

#### HDX-MS – Inhibitor study in pure particles

Continuous labelling, bottom-up hydrogen-deuterium exchange mass spectrometry was performed on SylP-reacted, ixazomib-bound and unbound $20S_{WT}$, respectively. All solutions used contained 1% DMSO (v/v), 100 mM NaCl and 50 mM $NaH_2PO_4$, pH 7.4. An initial equilibration step was carried out for each experiment, in which protein stocks were diluted to 6 µM into a $H_2O$-based buffer for a minimum of 30 min. Ixazomib was present at 10 µM in all solutions for the ixazomib-bound state. $D_2O$-based solutions containing identical additives to the equilibration buffers were adjusted to pD 7.4 using the standard electrode correction procedure (2). Protein stocks were diluted ten-fold into prepared $D_2O$-based solutions to initiate H/D exchange at room temperature. Aliquots were taken at 0.167 – 1440 minute time points from the solution undergoing exchange, and quenched by acidification to $pH_{read}$ 2.5 by mixing sample aliquots 1:1 (v/v) with 3 M guanidine hydrochloride, 250 mM $NaH_2PO_4$, pH 1.52, and 3 mM n-dodecylphosphocholine (3) and flash frozen in liquid nitrogen. Samples were stored at -80 °C prior to analysis. Undeuterated controls underwent identical sample preparation and handling, with the exception that all solutions were $H_2O$-based. See SI Appendix, Table. S1 for additional details.

An ACQUITY M-Class UPLC System equipped with HDX technology (Waters, Milford, MA, USA) was used for liquid handling and reverse-phase separation. A nepenthesin-2 column (Affipro, AP-PC-004, 1 mm × 20 mm, 16.2 µL) was used to digest 15 pmol of sample at 15 ºC. An ACQUITY BEH C18 (1.7µm, 2.1 mm × 5 mm; Part#: 186003975, Waters) Column then trapped the resulting peptides for 3 min at a flowrate of 100 µL/min before separating the peptides on an HSS T3 (1.8 µm, 1.0 × 50 mm; Part#: 186003535, Waters) column using a linear, $H_2O$: ACN gradient where both solutions were acidified with 0.1% formic acid. ACN was ramped from 5 – 35% over an 8-minute period using a flowrate of 100 µL/min at 0 ºC. To reduce column carry-over, the sample loop and protease column were cleaned between injections by flushing with 1.5 M guanidine hydrochloride, 4% (v/v) acetonitrile, 0.8% (v/v) formic acid, and 1.5 mM n-dodecylphosphocholine.

The UPLC outflow was directed to a SYNAPT G2-Si Q-TOF operated and calibrated, externally and dynamically, as described above. A range of 50 – 2000 m/z with a scan time of 0.4 sec was used to acquire data. Fragmentation was induced by linearly increasing the transfer collision energy in alternating scans over 20–40 V. To exclude smaller ions the quadrupole was manually set to dwell at 300 m/z. The TOF analyzer was operated in resolution mode. Ion mobility was controlled manually, and drift time-aligned $MS^E$ data-independent acquisition was employed for peptide mapping, both as described previously (4).

All reference and deuterated samples were repeated in technical triplicates. Peptide identification was carried out using $MS^E$ experiments of undeuterated samples searched against a database containing the plasmid sequences of both the α- and β-subunits in ProteinLynx Global Server (v3.0.3). Peptide filtering parameters were taken from Sørensen et al (5) and surviving spectra were then manually inspected to ensure sufficient signal-to-noise ratio using DynamX v3.0 (Waters). The significance interval for deuterium uptake was calculated using the statistical workflow described in (6). Data were visualized using heatmaps generated using HDgraphiX (7).

##### HDX-MS – Hybrid studies

Continuous labelling, bottom-up hydrogen-deuterium exchange mass spectrometry was performed on each of the seven hybrid particle populations, for both an apo and SylP-reacted state, with the exception of $20S_{T1A}$ due to it being incapable of binding ligand. All solutions used contained 100 mM KCl and 50 mM $KH_2PO_4$, pH 7.4. Initial protein stocks were prepared in $H_2O$-based buffer at 6 µM for 0, 28, 56, 81 and 100% $\beta_{WT}$ and 12 µM for 11% and 93% $\beta_{WT}$ before being stored at 0 °C. $D_2O$-based solutions containing identical additives to the equilibration buffers were adjusted to pD 7.4 using the standard electrode correction procedure (2). HDX labelling was carried out over 0.25 – 100 min using a PAL3 Robotic Tool Change System and Chronos Software (Leap Technologies/CTC Analytics). HDX was initiated by a 7.5-fold dilution into $D_2O$-based buffer at 20 ºC, resulting in 87% $D_2O$ (v/v) during labeling. Reactions were quenched by acidification to $pH_{read}$ 2.5 at 0 °C by mixing sample aliquots 1:1 (v/v) with 6 M guanidine hydrochloride, 250 mM $NaH_2PO_4$, pH 1.52, and 3 mM n-dodecylphosphocholine (3). See SI Appendix, Table. S2 for additional details.

An ACQUITY M-Class UPLC System equipped with HDX technology (Waters Corp) was used for reverse-phase separation. A nepenthesin-2 column (Affipro, AP-PC-004, 1 mm × 20 mm, 16.2 µL) was used to digest 20 or 40 pmol of sample at 15 °C prior to trapping for 3 min on a ACQUITY BEH C18 VanGuard precolumn (Waters Corp). Peptides were then separated on an ACQUITY BEH C18 column (Waters Corp) over a 12 min $H_2O$: ACN gradient at 0 ºC using a flowrate of 40 µL/min. ACN was initially ramped from 5 – 35% over 7 min before increasing to 85% over the next 30 sec and holding for 1.5 min. ACN was then dropped to 5% over the final 3 min.

Both solutions were acidified with 0.1% formic acid. Between samples, the sample loop and protease column were cleaned by flushing with 1.5 M guanidine hydrochloride, 4% (v/v) acetonitrile, 0.8% (v/v) formic acid, and 1.5 mM n-dodecylphosphocholine using the PAL3 Robotic Tool Change System.

The UPLC eluent was directed to a SELECT SERIES™ Cyclic™ IMS instrument operated in positive-ion mode, and in resolution V-mode. A range of m/z range of 250 – 2000 with a scan time of 0.4 sec was used to acquire data. The instrument was operated at a capillary voltage of 2.8 kV, cone voltage of 30 V, desolvation temperature of 175 °C, and source temperature of 80 °C. Peptides were identified using $HDMS^E$ acquisition in the absence of $D_2O$ with a transfer CE ramp from 20 to 29 V. Single-pass cIMS was used to separate overlapping peptides, with a 6 ms injection, 5 ms separation, and 17 ms ejection/acquire sequence.

All reference and deuterated samples were measured in technical triplicates. Peptide identification and filtering was done as stated above. The significance interval for deuterium uptake was calculated as stated above. To investigate cooperativity between hybridized subunits upon ligand binding, changes in the relative deuterium uptake values were plotted as a function of increasing SylP-reacted $\beta_{WT}$ population. The rate of change was calculated across all peptides and H/D exchange times to determine a global mean and standard deviation. Peptides showing changes greater than 2σ were deemed statistically significant. This analysis was then repeated to investigate changes as a function of neighbour identity, in which a separate global mean and standard deviation was calculated. Data were visualized using heatmaps as stated above, or as volcano plots generated using in-house python scripts.

#### SI Appendix Figures

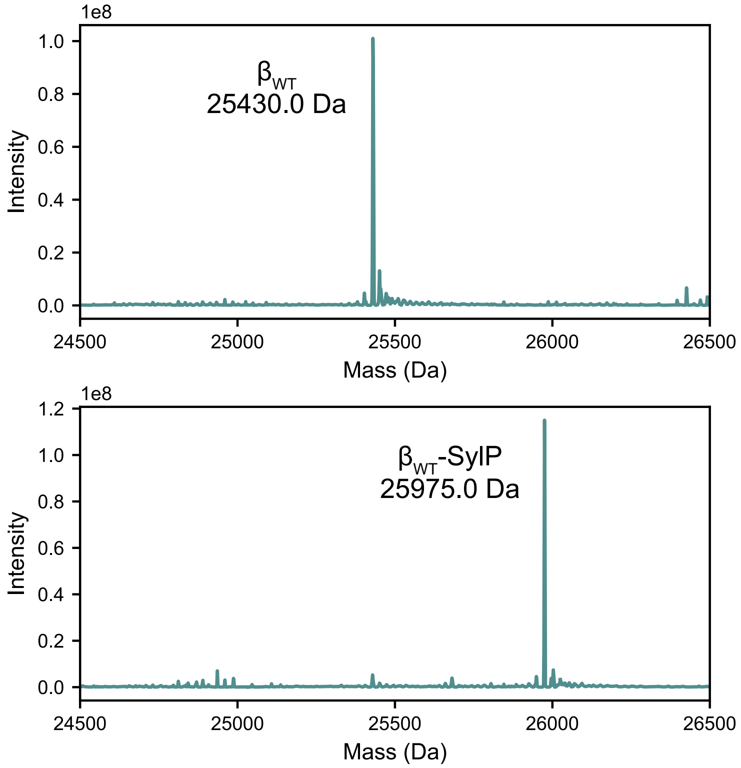


**Fig. S1.** Deconvoluted mass spectra of the apo (top) or SylP-reacted (bottom) $\beta_{WT}$-subunit reveal expected molecular weights. Spectra were summed over the chromatographic peak for the β-subunit prior to deconvolution. The measured molecular weights are labeled.

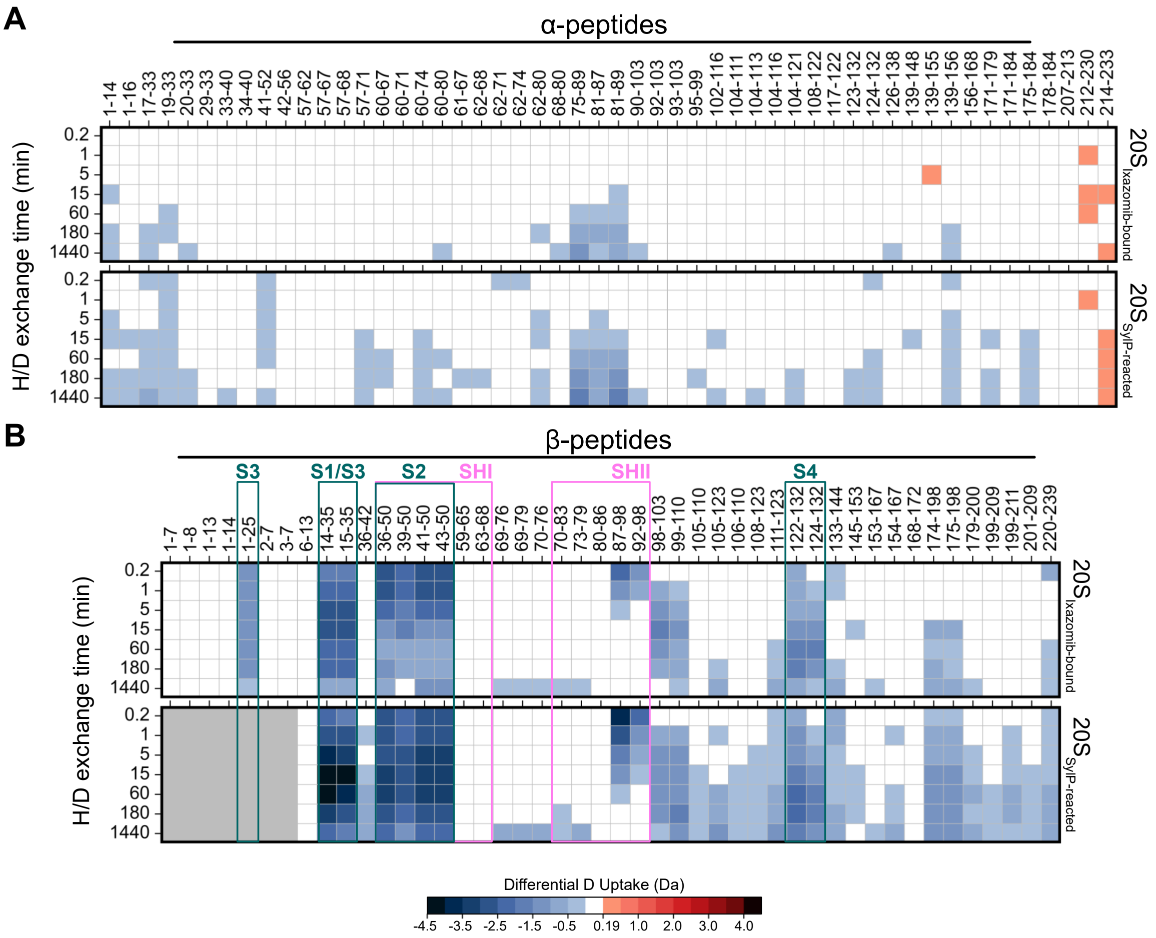


**Fig. S2.** Heatmap of the relative deuterium uptake in the α- and β-subunit in response to inhibitor binding, referenced to the apo $20S_{WT}$. Statistical analysis determined a significance interval of 0.19 Da which was used as a lower bound. All other changes were determined to be significant above 0.5 Da, with differences in deuterium uptake colour coded. Gray boxes indicate missing peptides due to the covalent modification by SylP. Peptides associated with the active site (S pockets) and switch helices are indicated.

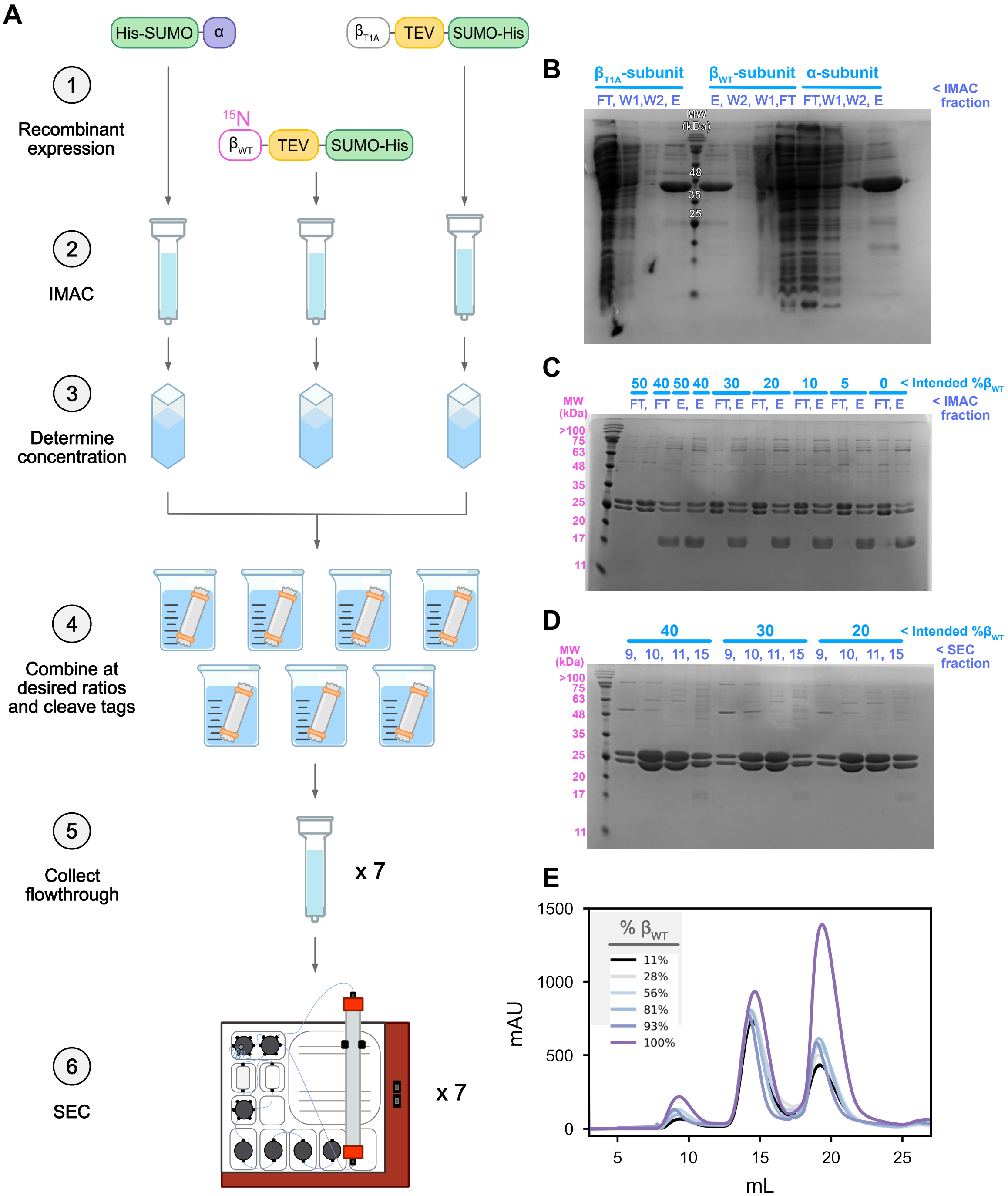


**Fig. S3**. Purification scheme to produce hybrid 20S CPs. Subunits are individually expressed and purified using IMAC (**A**, **B**). They are then combined at set ratios and allowed to oligomerize during tag-cleavage and dialysis. Cleaved tags are then removed via another round of IMAC (**C**) before the hybridized samples are placed on SEC to remove unassembled protomers (**D**, **E**).

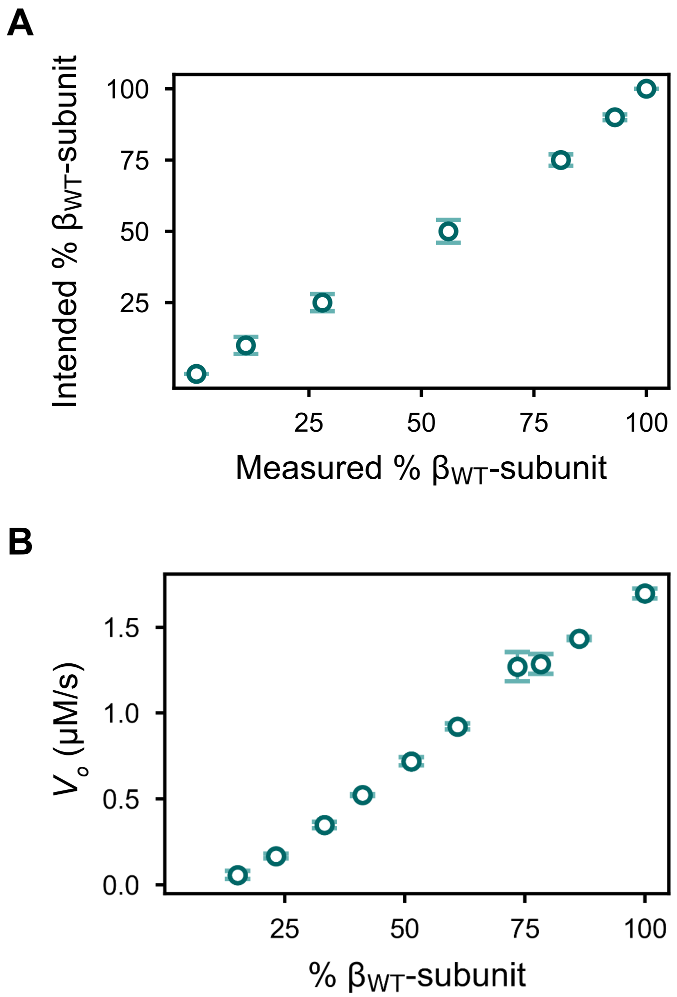


**Fig. S4**. Hybridized samples assemble to form functional core particles. (**A**) The measured ratio of $\beta_{WT}$: $\beta_{T1A}$ in pure hybridized samples, as determined via trypsin digest and MS analysis, are in good agreement with the intended ratios used to assemble the mixed particles during protein production. This suggests little to no partitioning between the subunit types during oligomerization; and (**B**) Catalytic activity of hybrid 20S CP samples against the tripeptide, Z-VLR-AMC. The data points and error bars represent averages and standard deviation (95% C.I.), respectively, calculated on the basis of three technical replicates.

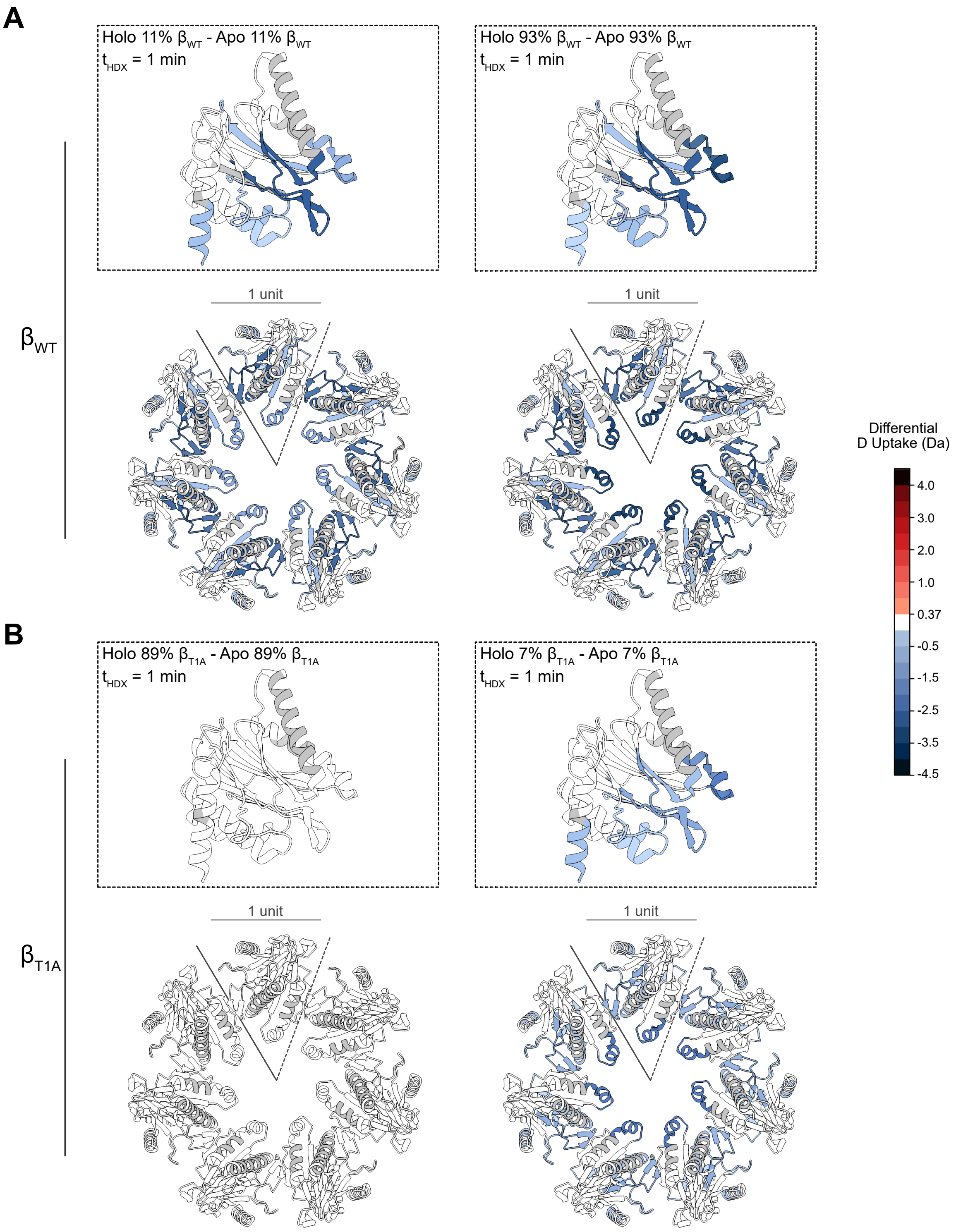


**Fig. S5.** Changes in the relative deuterium uptake of the β-subunits is dependent on the fraction of SylP-reacted $\beta_{WT}$-protomers present within the ring. (**A**) β-rings coloured according to the changes in the relative D-uptake at $t_{HDX}$ = 1 min of the $\beta_{WT}$-subunit upon SylP-reaction for the extreme ratios, 89:11 (left) and 7:93 (right) ($\beta_{T1A}$: $\beta_{WT}$) compared to their respective unbound controls; and (**B**) β-rings coloured according to the changes in the relative D-uptake at $t_{HDX}$ = 1 min of the $\beta_{T1A}$-subunit upon SylP-reaction for the extreme ratios, 89:11 (left) and 7:93 (right) ($\beta_{T1A}$: $\beta_{WT}$) compared to their respective unbound controls. Statistical analysis determined a significance interval of 0.19 Da which was used as a lower bound. All other changes were determined to be significant above 0.5 Da, with differences being coloured according to the colour bar. Uncovered regions are coloured gray. Structure visualized in Chimera 1.7 (PDB 9ce5).

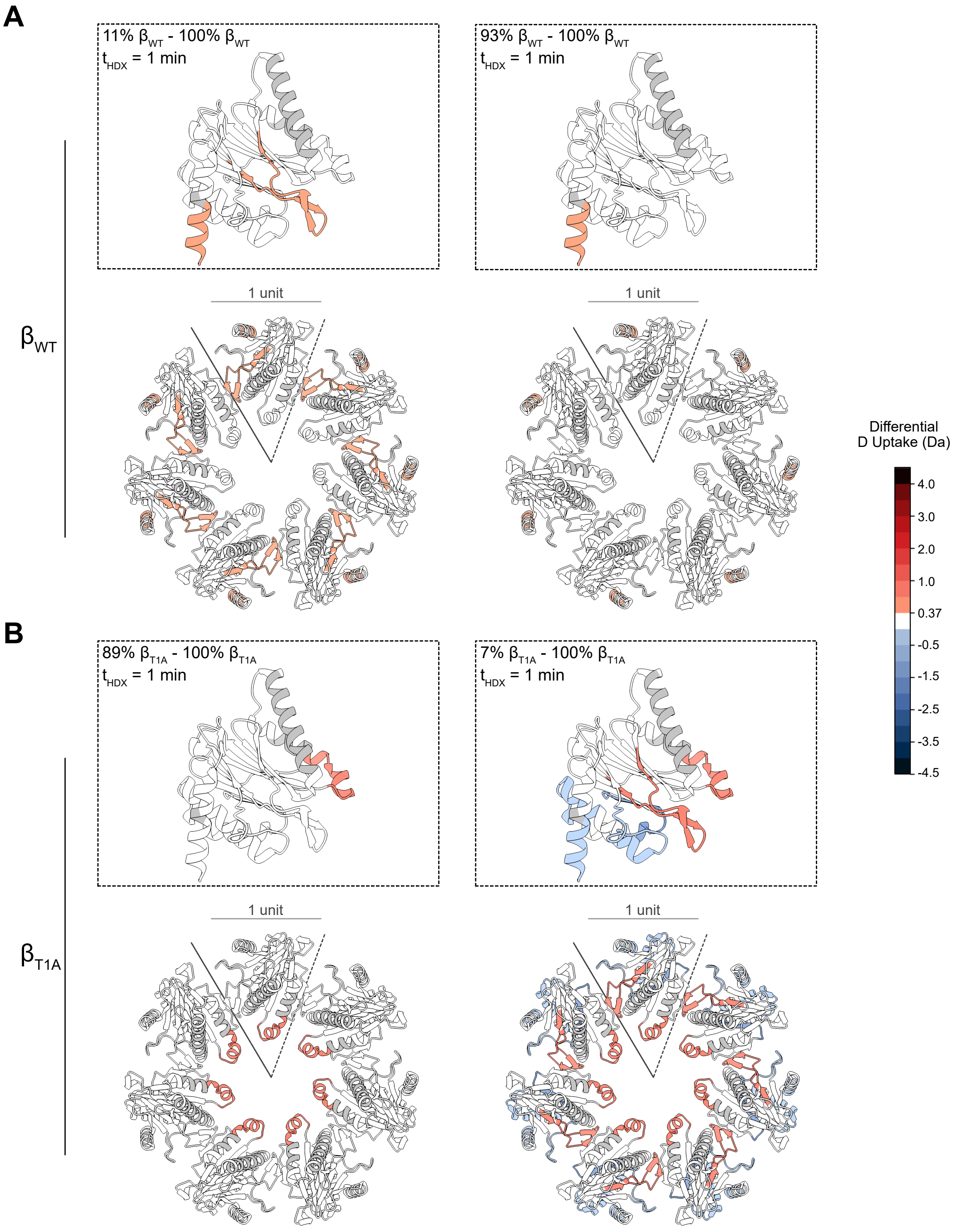


**Fig. S6**. Changes in the conformational dynamics of both β-subunit types are dependent on the fractional population of the opposing subunit. (**A**) β-rings coloured according to the changes in the relative D-uptake at $t_{HDX}$ = 1 min of the $\beta_{WT}$-subunit upon hybridization for the extreme ratios, 89:11 (left) and 7:93 (right) ($\beta_{T1A}$: $\beta_{WT}$) compared to the $20S_{WT}$ control; and (**B**) β-rings coloured according to the changes in the relative D-uptake at $t_{HDX}$ = 1 min of the $\beta_{T1A}$-subunit upon hybridization for the extreme ratios, 89:11 (left) and 7:93 (right) ($\beta_{T1A}$: $\beta_{WT}$) compared to the $20S_{T1A}$ control. Statistical analysis determined a significance interval of 0.19 Da which was used as a lower bound. All other changes were determined to be significant above 0.5 Da, with differences being coloured according to the colour bar. Uncovered regions are coloured gray. Structure visualized in Chimera 1.7 (PDB 9ce5).

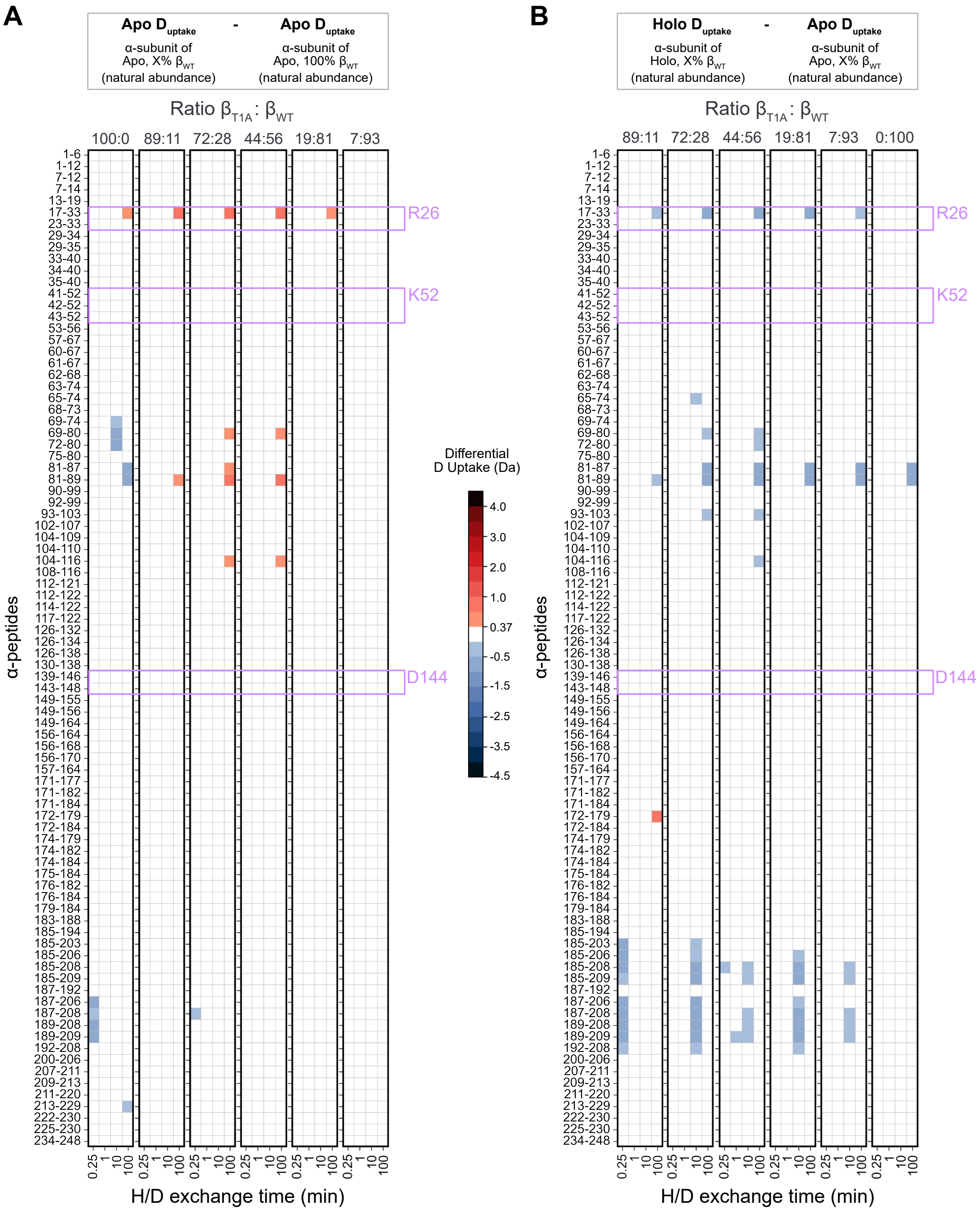


**Fig. S7**. The α-subunit allosterically senses changes within the β-rings. (**A**) Heatmaps depicting the relative deuterium uptake of the α-subunit within each ratio of hybridized core particle compared to that of a fully WT CP; and (**B**) Heatmaps depicting the relative deuterium uptake of the α-subunit of each SylP-reacted hybrid complexes compared to their unbound equivalent. Peptides containing residues important for the binding of regulatory particle are highlighted. Statistical analysis determined a significance interval of 0.19 Da which was used as a lower bound. All other changes were determined to be significant above 0.5 Da, with differences in deuterium uptake colour coded.

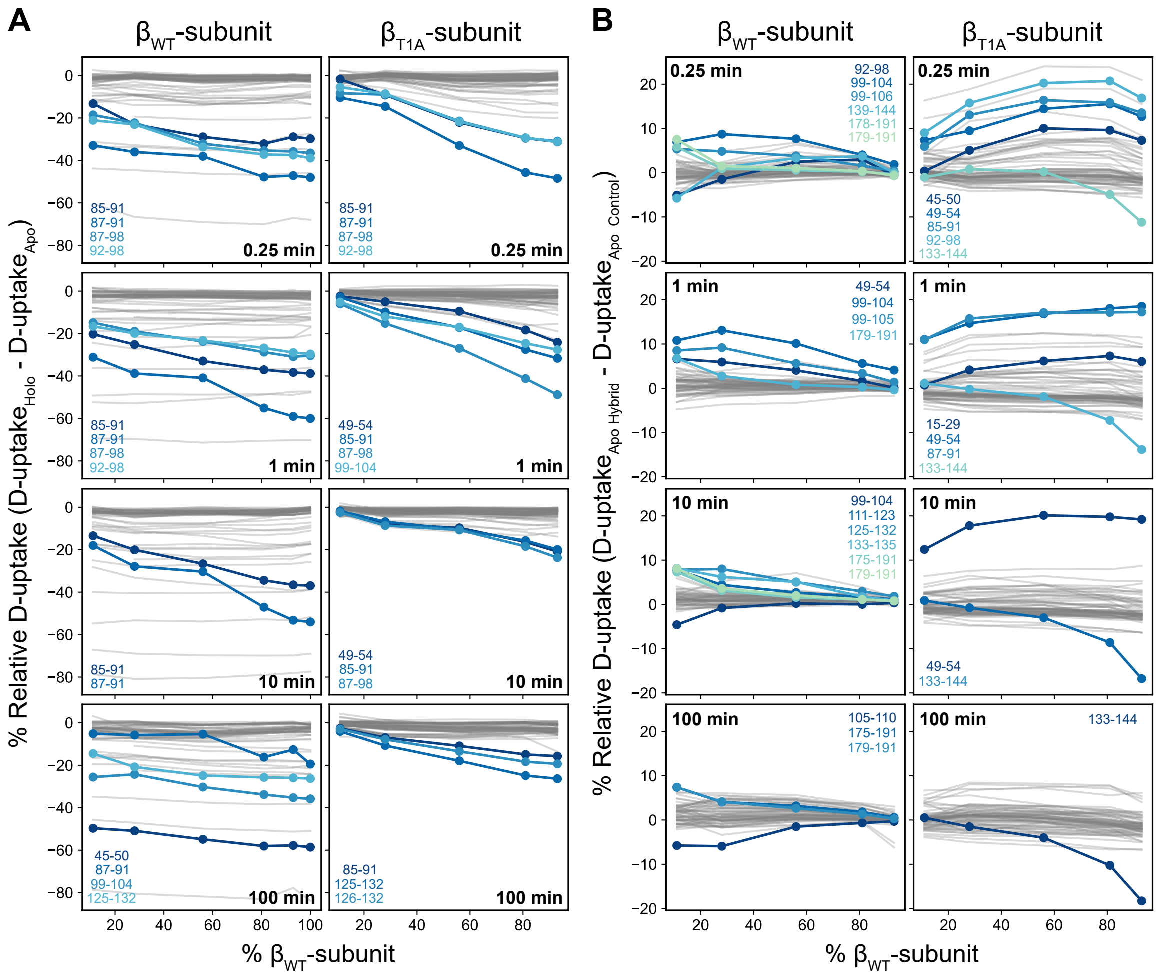


**Fig. S8**. Regions of allosteric communication can be identified through hybridization HDX-MS experiments. (**A**) Differences in the relative deuterium uptake of a given peptide between the holo and apo states of the hybrid complexes are plotted as a function of increasing population of SylP-reacted $\beta_{WT}$-subunit for both the WT and variant subunit at each H/D reaction time. This compositional slope was used to report on whether a peptide's conformational dynamics are coupled to ring composition in the ligand-bound state; and (**B**) Differences in a given peptide's relative deuterium uptake from both hybridized apo $\beta_{T1A}$- and $\beta_{WT}$-subunits compared to either the homogenous apo $20S_{T1A}$ or $20S_{WT}$, respectively, are plotted as a function of increasing unbound $\beta_{WT}$-subunit at each H/D reaction time. A global mean was calculated across each timepoint and subunit type for either (**A**) or (**B**), respectively, and peptides showing no significant change (within a 95% C.I.) relative to that mean are plotted as gray lines. Peptides showing significant change are coloured and labelled for each respective subunit type and reaction time.

**SI Appendix Tables**

**Table S1.** HDX summary table for HDX-MS of pure 20S CP.

| Dataset | Apo 20S $CP_{WT}$ | SylP-reacted 20S $CP_{WT}$ | Ixazomib-bound 20S $CP_{WT}$ |
|---|---|---|---|
| **HDX reaction details** | Final $D_2O$ concentration (v/v) = 90%<br>$pH_{corr}$ 7.4<br>Room temperature<br>1% DMSO (v/v) | Final $D_2O$ concentration (v/v) = 90%<br>$pH_{corr}$ 7.4<br>Room temperature<br>1% DMSO (v/v) | Final $D_2O$ concentration (v/v) = 90%<br>$pH_{corr}$ 7.4<br>Room temperature<br>10 µM Ixazomib<br>($K_D$ = 1.0 µM)<br>1% DMSO (v/v) |
| **HDX time course (min)** | 0.167, 1, 5, 15, 60, 180, 1440 | | |
| **HDX Undeuterated controls** | 3 | | |
| **Back-exchange** | α-subunit average back-exchange: 32.8% (range: 13.8 – 52.3 %)<br>β-subunit average back-exchange: 33.3% (range: 16.9 – 53.6 %) | | |
| **Number of peptides** | α-subunit: 52<br>β-subunit: 50 | α-subunit: 52<br>β-subunit: 43 | α-subunit: 52<br>β-subunit: 50 |
| **Sequence coverage** | α-subunit: 84%<br>β-subunit: 92% | α-subunit: 84%<br>β-subunit: 90% | α-subunit: 84%<br>β-subunit: 92% |
| **Average peptide length/redundancy** | α-subunit redundancy: 3.0<br>β-subunit redundancy: 2.6 | α-subunit redundancy: 3.0<br>β-subunit redundancy: 2.1 | α-subunit redundancy: 3.0<br>β-subunit redundancy: 2.6 |
| **Replicates** | Three technical replicates | | |
| **Repeatability** | ± 0.06 Da | | |
| **Significant differences in HDX** | 0.19 Da | | |

**Table S2.** HDX summary table for HDX-MS of hybrid 20S CP.

<table>
<tr><th>Dataset</th><th>Apo 20S $CP_{WT}$</th><th>Apo 20S $CP_{T1A}$</th><th>Apo Hybrid 20S (all ratios)</th><th>Holo Hybrid 20S (all ratios)</th></tr>
<tr><td>HDX reaction details</td><td colspan="4">Final $D_2O$ concentration (v/v) = 87%<br>$pH_{corr}$ 7.4; 20 °C</td></tr>
<tr><td>HDX time course (min)</td><td colspan="4">0.25, 1, 10, 100</td></tr>
<tr><td>HDX Undeuterated controls</td><td colspan="4">3</td></tr>
<tr><td>Back-exchange</td><td colspan="4">α-subunit average back-exchange: 32.8% (range: 13.8 – 52.3 %)<br>β-subunit average back-exchange: 33.3% (range: 16.9 – 53.6 %)</td></tr>
<tr><td>Number of peptides</td><td>α-subunit: 85<br>$\beta_{WT}$-subunit: 58</td><td>α-subunit: 85<br>$\beta_{T1A}$-subunit: 62</td><td>α-subunit: 85<br>$\beta_{WT}$-subunit: 58<br>$\beta_{T1A}$-subunit: 62<br><br>Note: the number of peptides for the $\beta_{WT}$-subunit is reduced to 55 for the 89:11 ($\beta_{T1A}$: $\beta_{WT}$) ratio</td><td>α-subunit: 85<br>$\beta_{WT}$-subunit: 55<br>$\beta_{T1A}$-subunit: 62<br><br>Note: the number of peptides for the $\beta_{WT}$-subunit is reduced to 52 for the 89:11 ($\beta_{T1A}$: $\beta_{WT}$) ratio</td></tr>
<tr><td>Sequence coverage</td><td>α-subunit: 98%<br>$\beta_{WT}$-subunit: 90%</td><td>α-subunit: 98%<br>$\beta_{T1A}$-subunit: 91%</td><td>α-subunit: 98%<br>$\beta_{WT}$-subunit: 90%<br>$\beta_{T1A}$-subunit: 91%</td><td>α-subunit: 98%<br>$\beta_{WT}$-subunit: 89%<br>$\beta_{T1A}$-subunit: 91%</td></tr>
<tr><td>Average peptide length/redundancy</td><td colspan="3">α-subunit redundancy: 3.6<br>$\beta_{WT}$-subunit redundancy: 3.1<br>$\beta_{T1A}$-subunit redundancy: 3.3<br><br>Note: $\beta_{WT}$-subunit redundancy dropped to 3.0 for the for the 89:11 ($\beta_{T1A}$: $\beta_{WT}$) ratio</td><td>α-subunit redundancy: 3.6<br>$\beta_{WT}$-subunit redundancy: 3.0<br>$\beta_{T1A}$-subunit redundancy: 3.3<br><br>Note: $\beta_{WT}$-subunit redundancy dropped to 2.8 for the for the 89:11 ($\beta_{T1A}$: $\beta_{WT}$) ratio</td></tr>
<tr><td>Replicates</td><td colspan="4">Three technical replicates</td></tr>
<tr><td>Repeatability</td><td colspan="4">± 0.04 Da</td></tr>
<tr><td>Significant differences in HDX</td><td colspan="4">0.37 Da</td></tr>
</table>
